# Modulation of voltage-dependent K^+^ conductances in photoreceptors trades off investment in contrast gain for bandwidth

**DOI:** 10.1101/344325

**Authors:** Francisco JH Heras, Mikko Vähäsöyrinki, Jeremy E Niven

**Affiliations:** Department of Zoology, University of Cambridge, Cambridge CB2 3EJ, UK; Sensapex Ltd, Oulu, Finland; School of Life Sciences, University of Sussex, Falmer, Brighton BN1 9QG, UK; Centre for Computational Neuroscience and Robotics, University of Sussex, Falmer, Brighton BN1 9QG, UK

**Author notes:** Correspondence /. Current address: Champalimaud Research, Champalimaud Center for the Unknown, 1400-038 Lisbon, Portugal (Francisco JH Heras).

## Abstract

Modulation is essential for adjusting neurons to prevailing conditions and differing demands. Yet understanding how modulators adjust neuronal properties to alter information processing remains unclear, as is the impact of neuromodulation on energy consumption. Here we combine two computational models, one Hodgkin-Huxley type and the other analytic, to investigate the effects of neuromodulation upon *Drosophila melanogaster* photoreceptors. Voltage-dependent K^+^ conductances: (i) activate upon depolarisation to reduce membrane resistance and adjust bandwidth to functional requirements; (ii) produce negative feedback to increase bandwidth in an energy efficient way; (iii) produce shunt-peaking thereby increasing the membrane gain bandwidth product; and (iv) inactivate to amplify low frequencies. Through their effects on the voltage-dependent K^+^ conductances, three modulators, serotonin, calmodulin and PIP2, trade-off contrast gain against membrane bandwidth. Serotonin shifts the photoreceptor performance towards higher contrast gains and lower membrane bandwidths, whereas PIP2 and calmodulin shift performance towards lower contrast gains and higher membrane bandwidths. These neuromodulators have little effect upon the overall energy consumed by photoreceptors, instead they redistribute the energy invested in gain versus bandwidth. This demonstrates how modulators can shift neuronal information processing within the limitations of biophysics and energy consumption.

## Introduction

The activity of neurons and neural circuits is modulated to adjust their properties to changes in state, affecting behaviour. Modulation can occur at large spatial scales, encompassing large numbers of neurons, or may be more localised, being restricted to small circuits or single neurons [1, 2, 3, 4, 5]. It can be brief or prolonged, and may be intrinsic or extrinsic to the circuits being modulated. A variety of substances from small gases (e.g. NO) to biogenic amines (e.g. serotonin, dopamine), peptides (e.g. FMRFamide) and other small molecules (e.g. ATP, PIP2) can act as modulators [2], altering their biophysical properties and, by doing so, affecting electrical signalling in neurons.

Many studies have focussed on the specific effects of modulators on voltage-dependent conductances, quantifying their effects on signalling and, in some cases, on behaviour [2]. For example, by acting on voltage-dependent conductances modulators can switch interneurons and motor neurons from following synaptic inputs directly to generating intrinsic bursting rhythms [6]. Studies of sensory neurons have also shown that modulators can alter their gain and frequency response (bandwidth), as well as increasing their coding precision [7]. But despite extensive characterisation of the effects of neuromodulators upon coding and behavioural consequences, their impact upon the energy consumption has been largely ignored.

Numerous lines of evidence suggest that energy consumption is an important factor both in the function and evolution of neurons, neural circuits and the nervous system [8, 9, 10]. Energy consumption can be considerable; the human brain is estimated to consume 20% of the basal metabolic rate, whilst the retina of the blowfly is estimated to consume 8% resting metabolic rate [8]. One of the main processes that consumes energy is the movement of ions across the bilipid membrane to support electrical signalling by graded and action potentials [11, 12, 13, 14] The properties of the protein channels through which ions move determine the preand postsynaptic ion flux across the bilipid membrane. For example, changes in the properties of the voltage-dependent ion channels that generate action potentials can alter their energy consumption by orders of magnitude [15, 11]. Ion flux is related to energy consumption primarily through the work done by the Na^+^ / K^+^ pump, which is powered by the hydrolysis of ATP molecules [16, 17]. Many neuromodulators alter ion flux by adjusting the properties of protein channels through which ions move. This provides a route by which neuromodulators can affect neuronal energy consumption and the energy efficiency of electrical signalling.

Here we assess this possibility in the R1-6 photoreceptors of the fruit fly, *Drosophila melanogaster*, using an analytic computational model of the photoreceptor to understand and interpret the biophysics accompanied by a Hodgkin-Huxley-type model to assess energy costs. The models incorporate the voltage-dependent K^+^ conductances expressed by these photoreceptors: a rapidly inactivating conductance encoded by the Shaker gene; a more slowly inactivating conductance encoded by the Shab gene; and, a non-inactivating conductance of unknown provenance (the ‘novel’ conductance) [18, 19]. Fruit fly photoreceptors, like those of other insects, encode information contained in light as a graded depolarisation. Consequently, voltage-dependent K^+^ conductances can change the filter properties of the membrane, altering the gain and bandwidth of electrical signals and have recently been shown to improve energy efficiency [20, 21, 22, 19, 23, 14]. The Shaker and Shab voltage-dependent K^+^ conductances of fruit fly photoreceptors are modulated by serotonin and PIP2 [24, 25, 26]. This provides an opportunity to characterise the changes in electrical signalling wrought by modulators in fruit fly photoreceptors, and determine whether such changes produce substantial effects upon energy consumption and efficiency.

## Materials and methods

### Hodgkin-Huxley characteristics of the K^+^ voltage-dependent conductances

We used a Hodgkin-Huxley (HH) formulation [27] to model the dynamics of the three voltage-dependent K^+^ conductances present in the fruit fly, *Drosophila melanogaster*, R1-6 photoreceptors. Activation and inactivation rates were obtained from previous studies [22, 25, 19]. In the following formulae describing conductance properties, steady-state activation and inactivation gating variables have no units, the voltage, *V*, is given in mV and the time constants are given in ms.

### Shaker

The fast-activating and fast-inactivating conductance encoded by *Shaker* has conductance *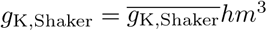*. The activation gating variable, *m*, has the following steady-state properties [25]

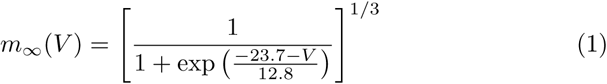

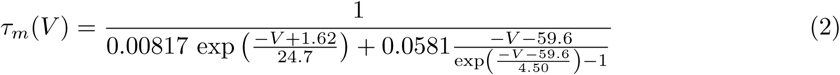

The inactivation gating variable *h* is the sum of two components, *h* = *ξ* + (1 *ξ*)*h*_1_. The first one is constant, representing the fraction of Shaker that fails to inactivate, which is *ξ* = 0.13. The second one is a HH variable with the following steady-state properties

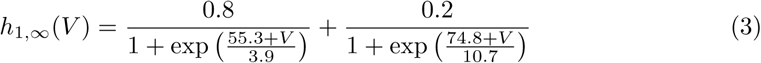

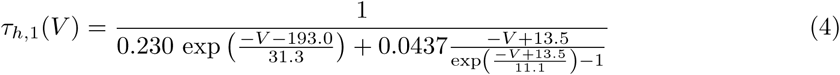

Serotonin produces a change in Shaker conductance properties (Fig 1). The activation time constant is shifted by 31.4 mV towards more positive voltages, while inactivation is shifted by 25 mV, with a small correction in shape [24]

**Figure 1:**
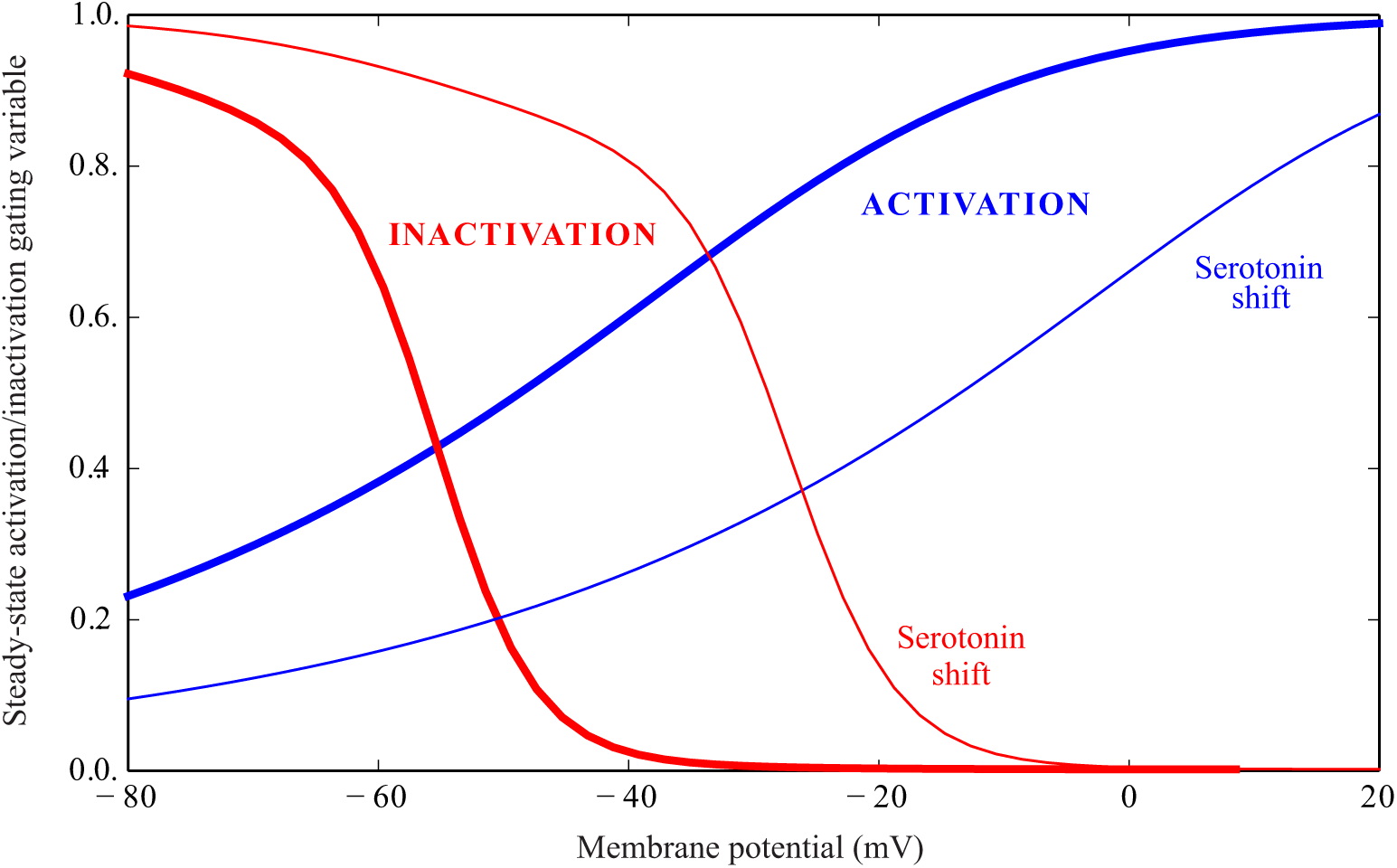
Steady-state activation and inactivation gating variables of Shaker in a *Drosophila* R1-6 photoreceptor. A fraction of the conductance, *ξ* = 0.13, fails to inactivate (not shown). Serotonin shifts Shaker activation and inactivation steady-state gating variables towards more depolarised potentials. The activation and inactivation curves are shown in bold blue and red lines, respectively. The shift in the activation and inactivation curves induced by serotonin is shown in thin lines.

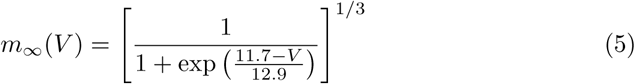

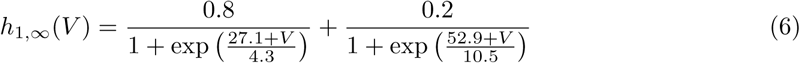

### Shab

The inactivating conductance encoded by *Shab* has conductance *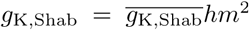*. The activation gating variable, *m*, has the following steady-state properties

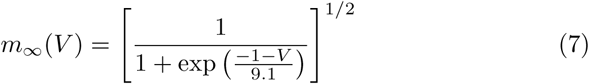

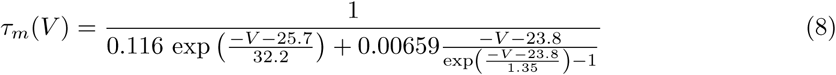

There are two modes of inactivation, with different kinetics, related to conformational changes produced when a K^+^ ion occupies the selectivity filter of the channel [28]. We modelled the inactivation as a sum of two variables representing each mode of inactivation *h* = *ξh*_1_ + (1– *ξ*)*h*_2_. The relative importance of the fast mode is considered to be constant around *ξ* = 0.7 [26].

We modelled both inactivation components with the same steady state

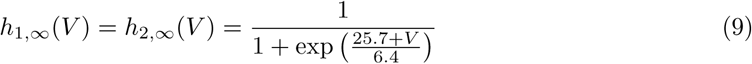

but different time constants

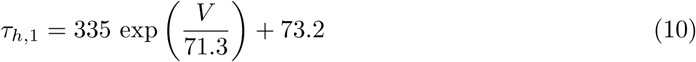

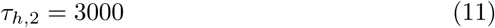

In the presence of serotonin, the steady-state curves and time constant curves are shifted towards positive potentials: activation by 13.7 mV and inactivation by 10 mV. In addition, the steady-state curves suffer small shape changes (Fig 2)

**Figure 2:**
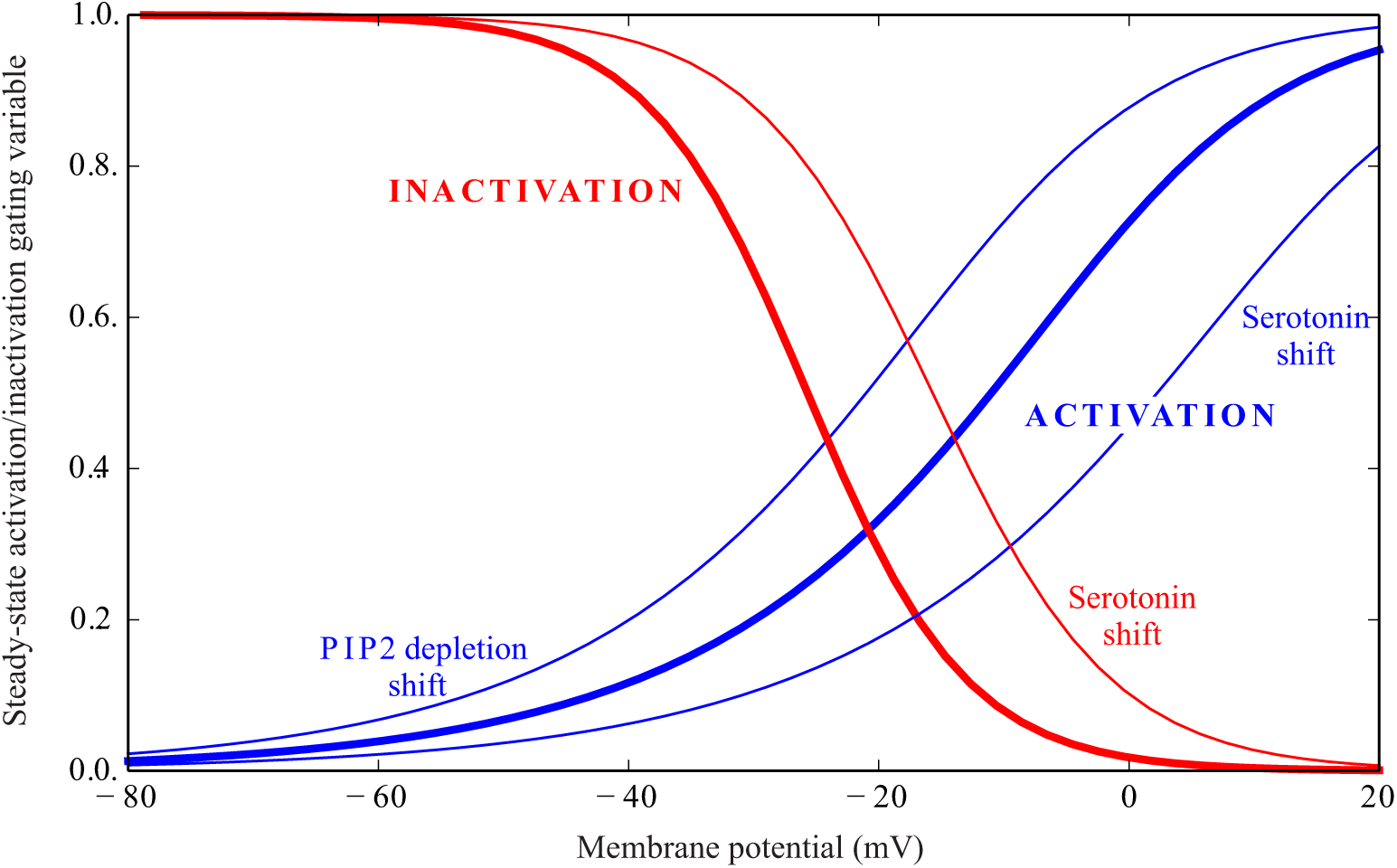
Steady-state activation and inactivation gating variables of Shab in a *Drosophila* R16 photoreceptor. Serotonin shifts Shab activation and inactivation steady-state gating variables towards more depolarised potentials, while PIP2 depletion by light shifts activation towards hyperpolarised potentials. The activation and inactivation curves are shown in bold blue and red lines, respectively. The shifts in the activation and inactivation curves induced by serotonin or PIP2 depletion are shown in thin lines.

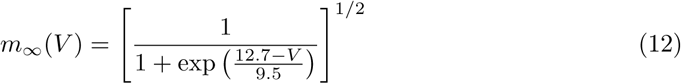

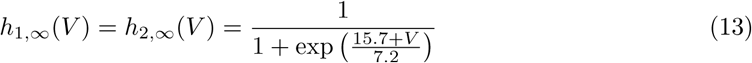

PIP2 depletion produces a leftward 10 mV shift in the Shab component of the conductance [26] (Fig 2). We model it as a 10 mV shift in the steady-state activation curve of Shab. Although [26] found some evidence that it affects more the slow inactivation component, we will apply the same shift to both components.

### Novel conductance

We modelled the non-inactivating K^+^ conductance discovered in double shaker,shab mutants [19], as *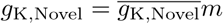*. The activation variable *m* has the following steady-state conductance and time constant

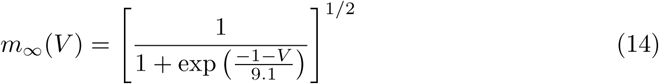

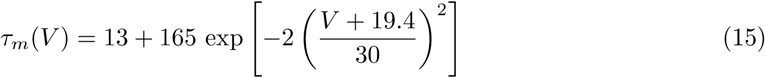

Steady-state activation variable, *m*_*∞*_(*V*), is shown in Fig 3.

**Figure 3:**
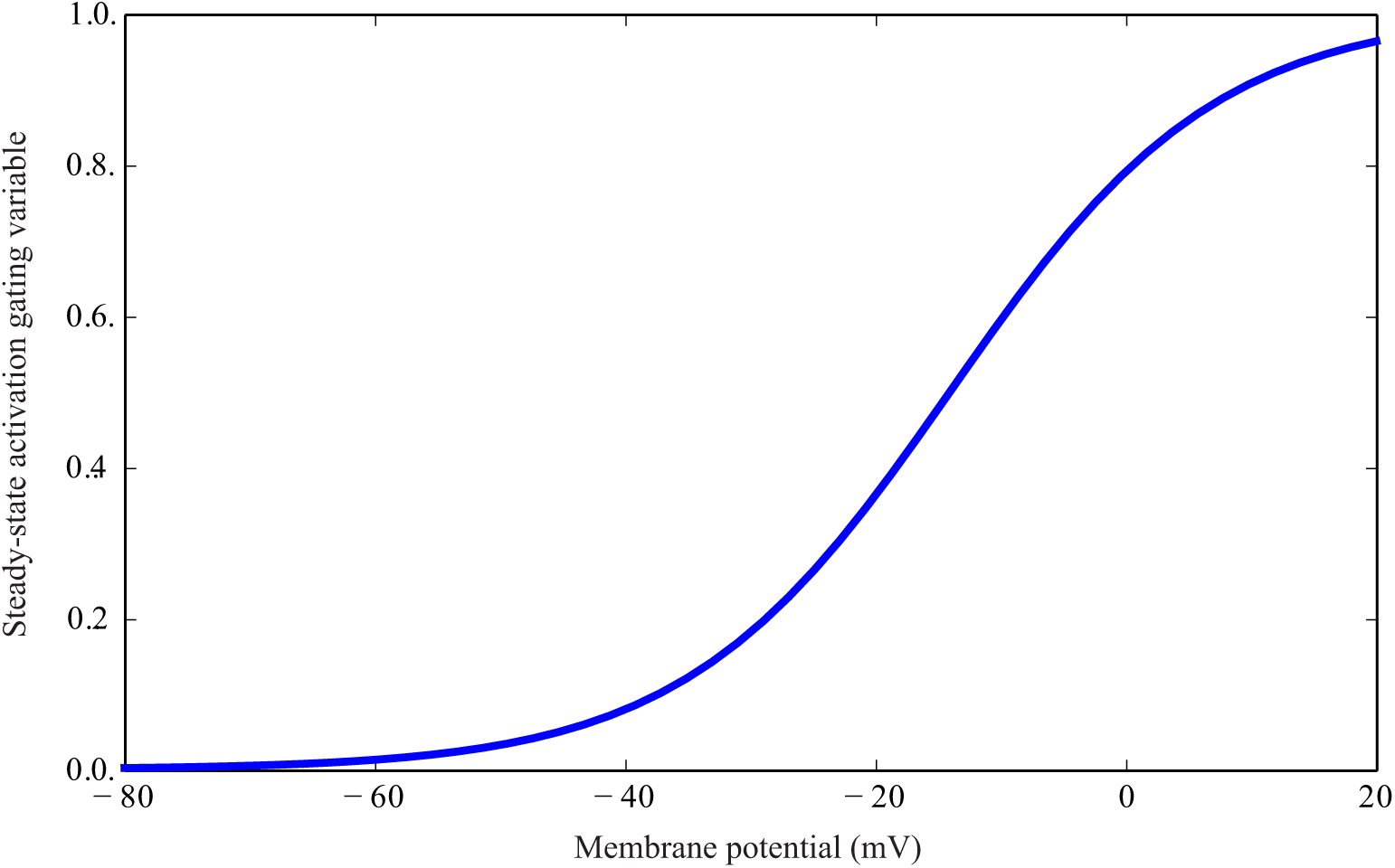
Steady-state activation gating variable of the Novel conductance in a *Drosophila* R1-6 photoreceptor.

### A biophysical model of the fruit fly R1-6 photoreceptor membrane

Our model of the fruit fly R1-6 photoreceptor membrane has capacitance 50 pF, intermediate among modern mesurements of 30 pF to 65 pF [25, 19]. It contains the three types of K^+^ conductances described above, namely Shaker, Shab and the Novel conductance, plus the light-induced curent (LiC). A Na^+^/K^+^ ATPase [29, 30] and a Na^+^/Ca^2^+ exchanger [31, 32, 33, 34, 35] maintain homeostasis by reversing the flux of ions. The K^+^ reversal potential, *E*_K_, is − 85 mV, and the reversal potential of LiC is *E*_L_ = 5 mV.

To reproduce a dark resting potential of −68 mV and input resistance in the dark of just below 300 MΩ (at the lower end of the *in vivo* electrophysiological experimental values, reported to be as low as a couple of hundred MΩ[36], 410 MΩ [22] or 1240 MΩ [37]) we included a 0.803 nS leak conductance with the reversal potential of the LiC and a 2.1 nS K^+^ leak conductance. From the dark resting potential, the photoreceptor model is depolarised by increasing the LiC. The recordings of the maximum steady-state depolarisation range from −45 mV [22] to almost −30 mV [37, 38], and we chose the intermediate value of −36 mV.

### Calculation of the steady state

We calculated the steady-state values of the conductances at different membrane potentials using well-known procedures. The total K^+^ conductance is the sum of the voltage-dependent K^+^ conductances and K^+^ leak conductance

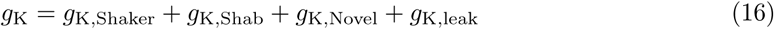

and the total depolarising conductance is the sum of the leak conductance and the light-induced conductance

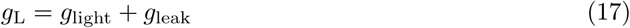

We assume that the LiC is mediated by Na^+^ (74%) and Ca^2+^ (26%), with a constant current fraction across voltages [34], and that only K^+^ flows throught the K^+^ conductances. At each steady-state depolarized membrane potential, *V*, we assume that all ionic currents, K+, Na^+^ and Ca^2^+, are restored by the Na^+^/K^+^ ATPase and the Na^+^/Ca^2+^ exchanger. In each cycle, the Na^+^/K^+^ ATPase pumps two K^+^ ions in and three Na^+^ ions out at the cost of one ATP molecule. The Na^+^/Ca^2+^ exchanger interchanges one Ca^2+^ by three Na^+^ ions. The net charge transported is 1 e per cycle in both the Na^+^/Ca^2+^ exchanger and the Na^+^/K^+^ ATPase.

This produces three equations, each equation being the result of zeroing one of the net ionic currents, *I*_(K)_, *I*_(Na)_ and *I*_(Ca)_

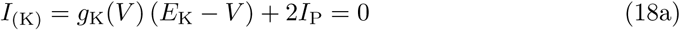

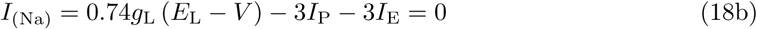

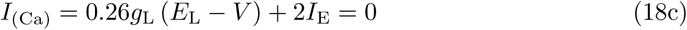

where *I*_P_ and *I*_E_ are the net currents produced by the Na^+^/K^+^ ATPase and the Na^+^/Ca^2+^ exchanger respectively. As we know the steady-state properties of the conductances, for each membrane voltage, *V*, we know *g*_K_(*V*) and we can solve the system of linear equations to obtain *ℊ* _light_, *I*_P_ and *I*_E_.

### Equivalent circuit for the calculation of the cell impedance

The response of a membrane containing voltage-dependent conductances to injected currents is, in the limit of small deflections, equivalent to the response of an electrical circuit composed of resistances, inductors and capacitors [27, 39, 40], sometimes called the equivalent RrLC circuit [41, 23, 14].

The equivalent circuit of an inactivating conductance, *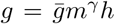*, has two phenomenological branches shunting (in parallel to) the membrane capacitance, *C*. The values of the electrical elements of the circuit, the resistances *R, r, r*_*h*_ and the impedances *L* and *L*_*h*_ are given by

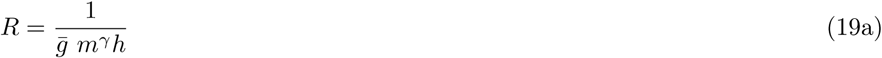

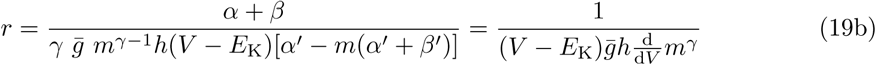

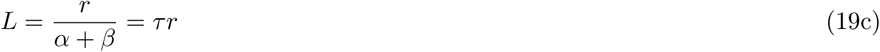

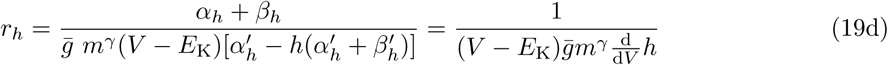

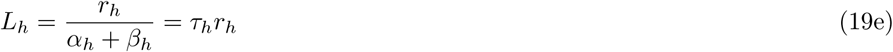

where *α* = *α*(*V*), *β* = *β*(*V*), *τ;* = *τ;* (*V*) are respectively the activation rate, deactivation rate and time constant of the HH activation variable, *m*, at the voltage, *V*. Similarly, *α*_*h*_ = *α*_*h*_(*V*), *β*_*h*_ = *β*_*h*_(*V*), *τ*_*h*_ = *τ*_*h*_(*V*) are respectively the activation rate, deactivation rate and time constant of HH inactivation variable, *h*, at the voltage, *V*. Primes (’) represent derivatives of the functions above with respect to voltage.

We then use the values of *R, r, L, r*_*h*_ and *L*_*h*_ to compute the impedance of the K^+^ conductance

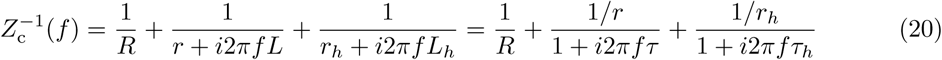

where *f* is the temporal frequency (in Hz) and *i* is the imaginary unit.

Please note that *R, r* and *L* are positive quantities, but *r*_*h*_ and *L*_*h*_ are negative quantities. *r*_*h*_ and *L*_*h*_ can be substituted by a positive resistance 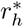 and positive capacitance 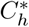 as described elsewhere [42]. As we only use the equivalent circuit to obtain the impedance, we do not perform this substitution.

If a conductance has different modes of inactivation or partial inactivation, the conductance can be divided into several conductances either non-inactivating or with only one mode of inactivation, and Eq 20 can be applied to each different component.

Once all the impedances of the different conductances, *Z*_*Shaker*_(*f*), *Z*_*Shab*_(*f*) and *Z*_Novel_(*f*), have been calculated using Eq 20, the impedance of the whole photoreceptor is

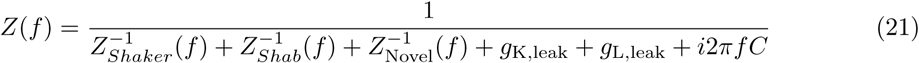

where *g*_K,leak_ and *g*_L,leak_ are the two leak conductances and *i*2*πf C* is the admittance of the membrane capacitance, *C*.

Note that for small deviations around the steady state, we consider *I*_P_ and *I*_E_ as fixed currents, which do not affect the photoreceptor impedance.

### Contrast gain

Each photon captured by the photoreceptor produces a transient increase in light-induced conductance and thus a transient increase in LiC (quantum bump). Thus, under physiological conditions, it is useful to describe how the photoreceptor voltage changes with changes in light level. A common measure is the contrast gain function (e.g. [43, 37]), which is the frequency dependent gain between light contrast and the voltage response of a photoreceptor. In this section we show that, under simplifying assumptions, the contrast gain is the product between LiC and impedance.

To account for the effect of voltage-dependent channels upon phototransduction gain, we implemented a simple transduction model. In this simple model, *g*_light_ follows light contrast, *c*(*t*), without any filtering or adaptation in the range of frequencies considered

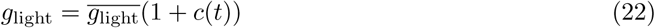

where *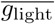* is the steady-state light induced conductance.

When the perturbations around the steady state, *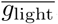*, are small, the resulting voltage fluctuations are much smaller than *E*_L_ –*V*, and the effect of the change in light contrast is equivalent to an injected current of zero mean

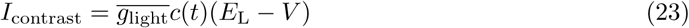

In the frequency domain, this current injected in a photoreceptor with membrane impedance *Z*(*f*) produces the following voltage fluctuations

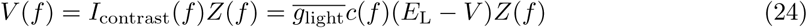

where, abusing the notation, we used the same symbol (*I*_contrast_(*f*) and *c*(*f*)) than in the time domain for the Fourier transforms. From this equation, we deduce that the contrast gain is

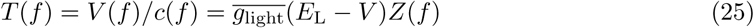

In a real photoreceptor, at low frequencies, contrast gain is reduced by adaptation. At high frequencies, light response is limited by the width and latency dispersion of the quantum bump [44, 45]. As a result, Eq 25 is a theoretical maximum, and will be a valid approximation only at intermediate frequencies.

### Energy consumption

The Na^+^/K^+^ pump uses energy from ATP hydrolysis to maintain ionic concentration gradients [17, 29]. This is the dominant cost of the photoreceptor [46, 14].

We obtain the net pump current solving Eqs 18. Because the Na^+^/K^+^ ATPase hydrolyses an ATP molecule for every 2 K^+^ ions pumped in and 3 Na^+^ ions pumped out [29], we can calculate the pump energy consumption as *|I*_*P*_ *| /e*. Units are hydrolysed ATP molecules/s when the current, *I*_P_, is given in amperes and the elementary charge, ***e*,** is in coulombs.

## Results

We modelled the electrical properties of a fruit fly R1-6 photoreceptor membrane to explore the effects of a combination of voltage-dependent K^+^ conductances on both the membrane filter properties and the photoreceptor energy consumption. We modelled the photoreceptor soma as a single electrical compartment, ignoring the effect of the axon. We implemented three types of voltage-dependent K^+^ conductances: a rapidly activating and inactivating current encoded by *Shaker* [18], a slowly activating and inactivating current encoded by *Shab* [18, 19] and a very slowly activating/non-inactivating current, the “Novel” conductance [19]. We used a Hodgkin-Huxley (HH) framework [27] for the K^+^ conductances, with activation and inactivation rates obtained from previous studies [22, 25, 19].

### Negative and positive feedback shape the voltage responses to injected current

The electrical properties of the membrane, including the voltage-dependent activation and inactivation of conductances, determine the voltage response to injected current. We simulated the voltage responses of the photoreceptor to small injected current pulses to separate the contribution of the different voltage-dependent conductances (Fig 4). We performed these simulations at different light depolarisations, obtained by increasing the light-induced conductance.

**Figure 4:**
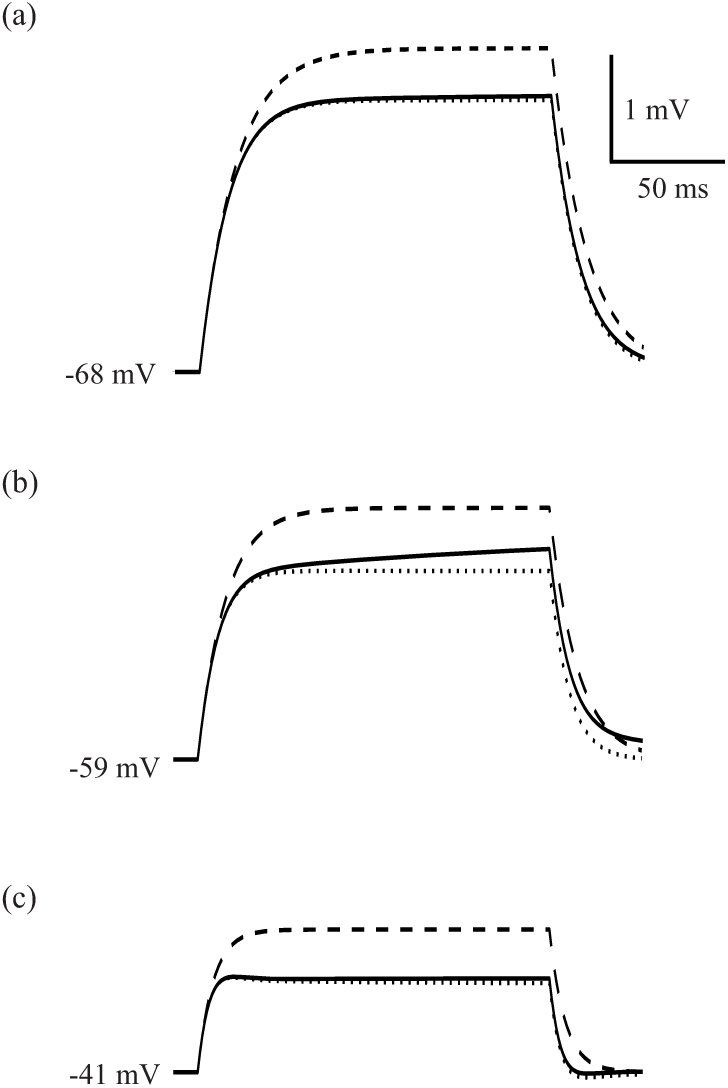
Voltage responses of simulated *Drosophila melanogaster* R1-6 photoreceptors to 0.01 nA injected current pulses (a) Voltage response of a dark adapted membrane. Three curves are plotted. Continuous line shows the result of a simulation where all gating variables are allowed to change following Hodkgin-Huxley equations, and thus the voltage response shows both negative and positive feedback. Dotted line is the result of a simulation where inactivation gating variables of all conductances are kept fixed in their steady-state value, and thus there is negative feedback but no positive feedback. Dashed line shows the result of keeping all gating variables (activation and inactivation) fixed during simulation, or equivalently, the voltage response of a passive membrane with the same membrane resistance and capacitance. (b) Same as (a) but in a photoreceptor that has been depolarised by light to −59 mV. (c) Same as (a) and (b) but in a photoreceptor that has been depolarised by light to −41 mV.

At all light depolarisations, activation of a voltage-dependent K^+^ conductance performs negative feedback, while inactivation of a voltage-dependent K^+^ conductance performs positive feedback. To distinguish the effects of negative and positive feedback, we freeze —i.e. we selectively keep fixed in their steady-state values— the activation and/or inactivation gating variables. When the inactivation gating variables are frozen positive feedback disappears. When all gating variables are frozen both positive and negative feedback disappear, and the voltage response reverts to the RC charging curve of a passive membrane.

The voltage responses of photoreceptors with active membranes (henceforth referred to as active photoreceptors) initially follow the RC charging curve of the passive photoreceptor (Fig 4) but diverge after a few milliseconds because the negative feedback produced by the activation of voltage-dependent conductances reduces the voltage deflection. Conductance inactivation is slower than activation, so the positive feedback produced from inactivation appears later than the negative feedback produced by activation.

The dark adapted photoreceptor produces the largest voltage responses to injected current (Fig 4a) because there is little activation of voltage-dependent conductances at −68 mV. There is, however, some negative feedback due to the fast activation of Shaker and the comparatively slower activation of Shab, reducing the voltage signal of the active photoreceptor below that of the passive photoreceptor by about 15%. After a couple of tens of milliseconds, Shaker inactivation produces a small amplification (approximately 1.5% after 150 ms).

Increasing the light induced conductance, as occurs in low light intensities, depolarizes the simulated photoreceptor to –59 mV. The membrane resistance decreases because of the increased light-induced and K^+^ conductances. As a consequence, the voltage responses become smaller (Fig 4b). Negative feedback becomes more apparent, subtracting approximately 25% from the voltage response, whilst positive feedback due to Shaker fast inactivation amplifies the depolarisation by approximately 9% at the end of a 150 ms pulse (Fig 4b), and up to 15% in longer pulses (not shown).

At high light levels, when the light-induced conductance depolarizes the model photoreceptor to –41 mV, the membrane resistance is further decreased and the voltage responses become smaller (Fig 4c). Negative feedback due to conductance activation becomes conspicuous, decreasing gain by approximately 37%, thereby producing an overshoot and an after-potential. Positive feedback due to conductance inactivation is reduced (less than 4% after a 150 ms pulse). At the end of the 150 ms pulse, about 20% of the positive feedback is due to Shab inactivation (not shown). This positive feedback is more apparent in longer time scales because of the slow dynamics of Shab inactivation, when it is responsible of about one third of the total inactivation (not shown).

### Negative and positive feedback shape the membrane impedance

To analyse the contribution of the voltage-dependent conductances to the membrane’s frequency response, we calculated the impedance, *Z*(*f*), which determines the linear filtering characteristics of the membrane in the frequency domain. By solving an equivalent electrical circuit that represents the linearization of the membrane incorporating voltage-dependent conductances, we obtained Eq.21 to calculate the impedance (see Material and methods) [27, 39, 40, 23, 14]. Using Eq.21, we calculated the impedance at each light level, depolarising the photoreceptor by increasing the light conductance.

In the dark, the membrane is a low-pass filter (Fig 5a), not very different from an RC filter with the same resistance and impedance. The feedback produced by voltage-dependent conductances lowers the impedance of the active membrane at every frequency to below that of the passive membrane with the same membrane resistance and capacitance. At low light levels, when the photoreceptor is depolarized to between –59 mV and –50 mV, the impedance shows a conspicuous low frequency amplification below 2-3 Hz (Fig 5a) that corresponds to the positive feedback due to Shaker inactivation. This amplification is produced solely by positive feedback from Shaker inactivation, and does not depend solely upon a window current as previously suggested [22]. At high light levels the photoreceptor is depolarized to above –41 mV, the negative feedback causes the impedance to become band-passing, and the Shaker-mediated amplification disappears (Fig 5a). Only at the highest light levels does the positive feedback from Shab inactivation amplify frequencies below 1 Hz. At medium and high light levels, the Novel conductance produces negative feedback that decreases impedance at low frequencies diminishing the low frequency amplification of Shab.

**Figure 5:**
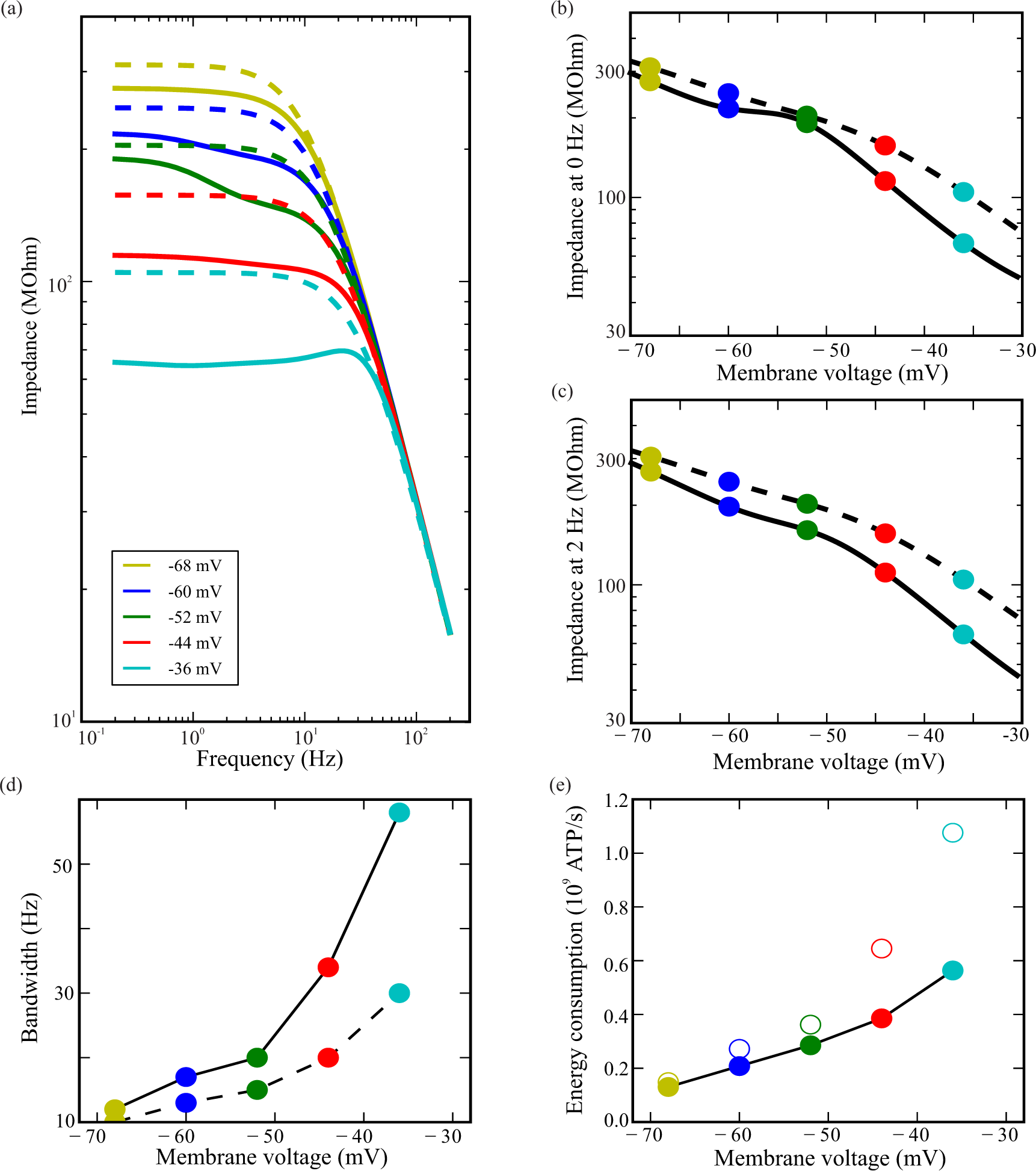
The effects of feedback produced by K^+^ conductances on a model photoreceptor, at dark resting potential and depolarised by light-induced current to four steady-state membrane potentials Photoreceptor impedance in simulated photoreceptors depolarised by light to different voltages (solid lines). Dashed lines represent the membrane impedances of photoreceptors with frozen conductances, i.e. where the activation and inactivation gating variables are kept constant at each steady state when computing the impedance. They are RC membranes with the photoreceptor capacitance and true membrane capacitance fixed at the value taken in that particular voltage. The low frequency limits and values of the impedance at 2 Hz are represented in subfigures (b) and (c). (d) The bandwidth of simulated photoreceptors with active K^+^ conductances (solid line) depolarised by light to different voltages versus the bandwidth of photoreceptors with frozen conductances at each voltage (dashed line). (e) Photoreceptor energy consumption increases with depolarisation. At each depolarisation, the cost of an active fruit-fly photoreceptor (filled circles) is smaller than that of a passive photoreceptor with the same bandwidth and capacitance (empty circles).

### Negative feedback reduces the cost of membrane bandwidth and increases the gain-bandwidth product

Many of the mechanisms that adapt photoreceptors to the prevailing light conditions, such as Ca^2+^ inhibiting TRP and TRPL channels [47] and closing the intracellular pupil [48], would attenuate low frequencies. The attenuation of low frequencies in the LiC affects the overall photoreceptor frequency response, masking the amplification that is seen in the membrane. Consequently, there is no significant linear amplification in the light response [49]. This suggests that amplification below 2 Hz is unlikely to be the primary function of the Shaker conductance and is merely a side effect of the relatively slow inactivation of fruit fly voltage-dependent conductances. Thus, when calculating gain and bandwidth we only consider frequencies above 2 Hz. The photoreceptor membrane increases its bandwidth from around 26 Hz in the dark adapted state to more than 100 Hz at the highest light levels (Fig 5d).

Photoreceptor energy consumption at each depolarisation level can be calculated from the flux of Na^+^ and K^+^ ions across the membrane, which must be pumped back across to the opposite side by the Na^+^/K^+^ pump (see Methods). The 4-fold increase in photoreceptor bandwidth from the lowest to the highest light levels is produced at the cost of a 4.3-fold increase in energy consumption, from 1.3× 10^8^ ATP*/*s in the dark to 5.6 × 10^8^ ATP*/*s at the highest light levels (Fig 5e).

At all light levels, the conductance properties boost the bandwidth of the photoreceptor membrane above that of a passive membrane with identical resistance and capacitance (Fig 5d). In the dark, the active photoreceptor membrane has a 6 Hz higher bandwidth than the corresponding passive membrane, increasing to 10 Hz at low light levels. At the highest light levels, the active photoreceptor membrane has a 40 Hz higher bandwidth than the corresponding passive membrane, an improvement of more than 70%. To achieve the same bandwidth as the active photoreceptor membrane with a passive membrane would increase the energy cost by 10% at low light levels rising to 40% at high light levels (Fig 5e).

### Negative feedback increases the gain-bandwidth product (GBWP)

Using the active and passive membrane impedances, we calculated the gain-bandwidth product (GBWP), the product of bandwidth and peak impedance [23]. The GBWP of the active photoreceptor membrane generally increases with depolarisation, though there is a pronounced ‘dip’ at intermediate voltages around −52 mV (Fig 6). In contrast, the GBWP of the corresponding passive membranes remains constant.

**Figure 6:**
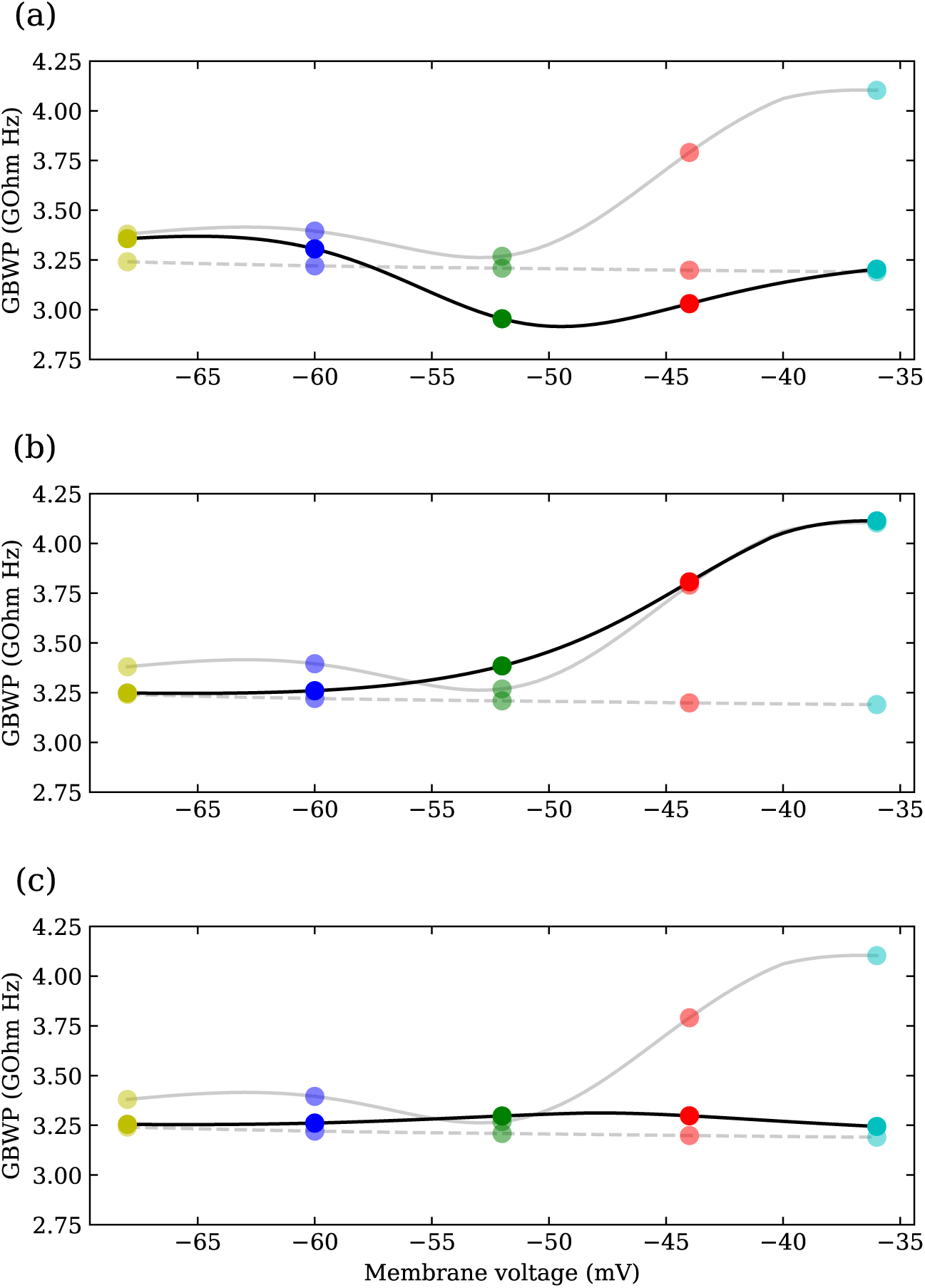
The contributions of the three voltage-dependent K^+^ conductances to the gainbandwidth product (GBWP) of the photoreceptor membrane, over the range of depolarisation produced by the light induced current. (a) GBWP of a fruit fly model photoreceptor across different light levels. Circles joined with black solid lines result from a model where only Shaker activation and inactivation gating variables are active, while the other two conductances are frozen, i.e. kept constant at each steady state when computing the impedance. Circles joined with grey solid lines are the result of the model with all active conductances, and circles joined with dashed lines are GBWP of the passive membrane, i.e. when all gating variables are frozen. (b) Same as (a) but keeping Shab activation and inactivation gating variables active, and freezing those of Shaker and Novel. (c) Same as (a) but keeping the Novel activation gating variable active, and freezing those of Shaker and Novel.

The increase in GBWP in the active photoreceptor is produced by the voltage-dependent K^+^ conductances. Changes in the membrane potential cause the voltage-dependent ion channels to alter their conformation changing their conductance after a delay, causing them to act like an inductance shunting (in parallel to) the membrane capacitance. This is akin to shunt peaking in an electronic amplifier, a technique in which an inductor is placed in series with the load resistance to compensate for capacitive effects and improve the frequency response [50, 51]. In the photoreceptor membrane the inductive effects of voltage-dependent K^+^ conductances increase the GBWP [23]. At low light levels shunt peaking increases GBWP by 10-15%, and by 25% at the highest light levels.

By selectively freezing the gating variables of the different voltage-dependent conductances, we distinguished the contributions of each conductance to shunt peaking (Fig 6). Shaker activation is the major contributor to shunt peaking at low light levels (membrane voltages below –55 mV). At higher light levels, however, Shab activation becomes the major contributor to shunt peaking. Activation of the Novel conductance is too slow to significantly contribute to shunt peaking at any light level.

### Inactivation of voltage-dependent K^+^ conductances affects photoreceptor bandwidth and energy consumption

The voltage-dependent Shaker and, to a lesser extent, the Shab conductances in *Drosophila* R1-6 photoreceptors inactivate. To assess the role of inactivation in these voltage-dependent K^+^ conductances, we froze the inactivation variable at different light levels, and assessed the bandwidth and energy consumption of these photoreceptor membranes across the full operating range of the photoreceptor (Fig 7). All other features of the photoreceptors remained identical. At low light levels, close to the resting potential there is little inactivation of the voltage-dependent K^+^ conductances. Membranes with inactivation frozen at these low light levels have a similar bandwidth and energy consumption to typical photoreceptors. As light levels increase and the photoreceptor depolarises, however, the bandwidth and energy consumption of these membranes is substantially higher than that of a typical photoreceptor (Fig 7a and b).

**Figure 7:**
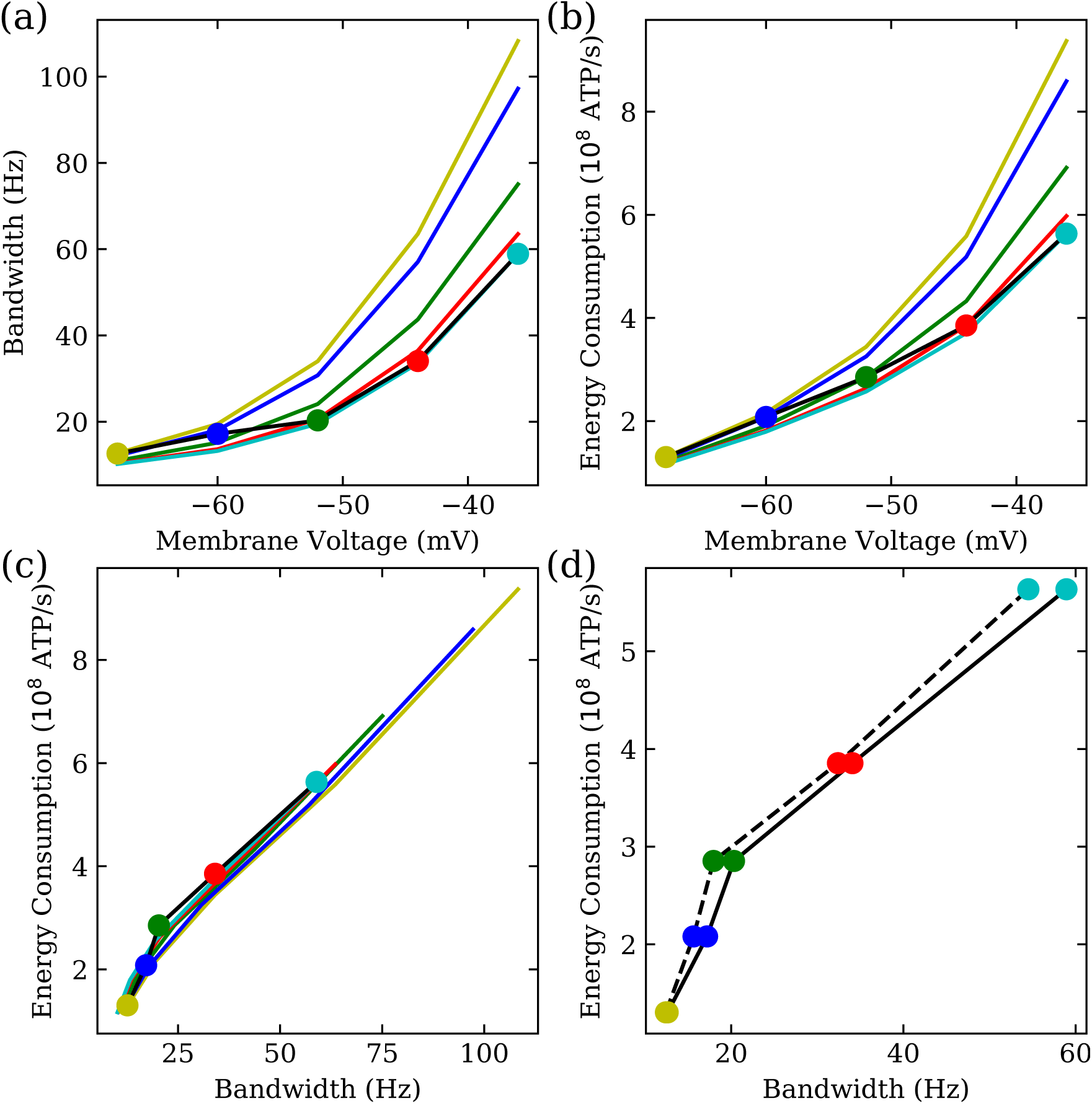
Inactivation of the voltage-dependent K^+^ conductances affects the bandwidth and energy consumption of photoreceptors. (a) Bandwidth at different membrane voltages of the photoreceptor (black line with colour circles) compared to that of photoreceptors where inactivation of the voltage-dependent conductances has been frozen, i.e. kept constant at the value corresponding to the circle of the same color. (b) As in (a), but plotting energy consumption against membrane voltage. (c) As in (a), but plotting energy consumption against bandwidth. (d) As in (c), but now the intact model (continuous line) is compared to a model in which the dynamics of inactivation have been accelerated so as to be the same as those of activation (dashed line).

When inactivation is frozen at high membrane potentials that correspond to high light levels, photoreceptor bandwidth and energy consumption is unaffected at high and at mid light levels (Fig 7a and b). Conversely, at low light levels these membranes have a lower energy consumption but also a lower bandwidth. Although the increase in bandwidth at low light levels is small in absolute terms, it is relatively large. There is almost no change in the energy cost of bandwidth itself, so energy savings are made not through increased energy efficiency but rather by reducing bandwidth at high light levels (Fig 7c). Thus, inactivation of a voltage-dependent K^+^ conductance permits relatively high bandwidths and energy costs at low light levels but ensures that there is no corresponding increase in bandwidth and energy costs at high light levels.

As an alternative to inactivating voltage-dependent K^+^ conductances, the excessive energy consumption and bandwidth at depolarised potentials could be reduced using non-inactivating conductances with the same steady-state parameters. To study this case, we accelerated the dynamics of inactivation of both Shaker and Shab voltage-dependent K^+^ conductances to be identical to those of activation. We found that the energy consumption of bandwidth is increased at all depolarised membrane potentials (Fig 7d). This reduction in the energy efficiency of the modified model is a consequence of decreased negative feedback caused by the weaker activation of the voltage-dependent conductances. Thus, the inactivation voltage-dependent K^+^ conductances improves the energy efficiency of photoreceptors, reducing the energy cost of bandwidth.

### Feedback decreases contrast gain and decreases the cost of phototransduction gain-bandwidth product

Photoreceptor contrast gain depends not only on the membrane impedance but also on the light-induced current (LiC). To determine how the membrane filter properties affect photoreceptor contrast gain, we incorporated a simplified, infinitely fast phototransduction cascade (Eqs 22, 23). Under these simplifying conditions, the contrast gain is the product of the LiC and membrane impedance (Eq 25).

Feedback from the voltage-dependent K^+^ conductances changes the impedance but does not change the LiC. Consequently, negative feedback from activation decreases the contrast gain of an active photoreceptor compared to that of a passive photoreceptor with identical membrane resistance and capacitance (Fig 8a). The net consequence is that, in our model, voltage-dependent conductances decrease the photoreceptor peak contrast gain (i.e. the maximum of contrast gain across different frequencies) at all light intensities.

**Figure 8:**
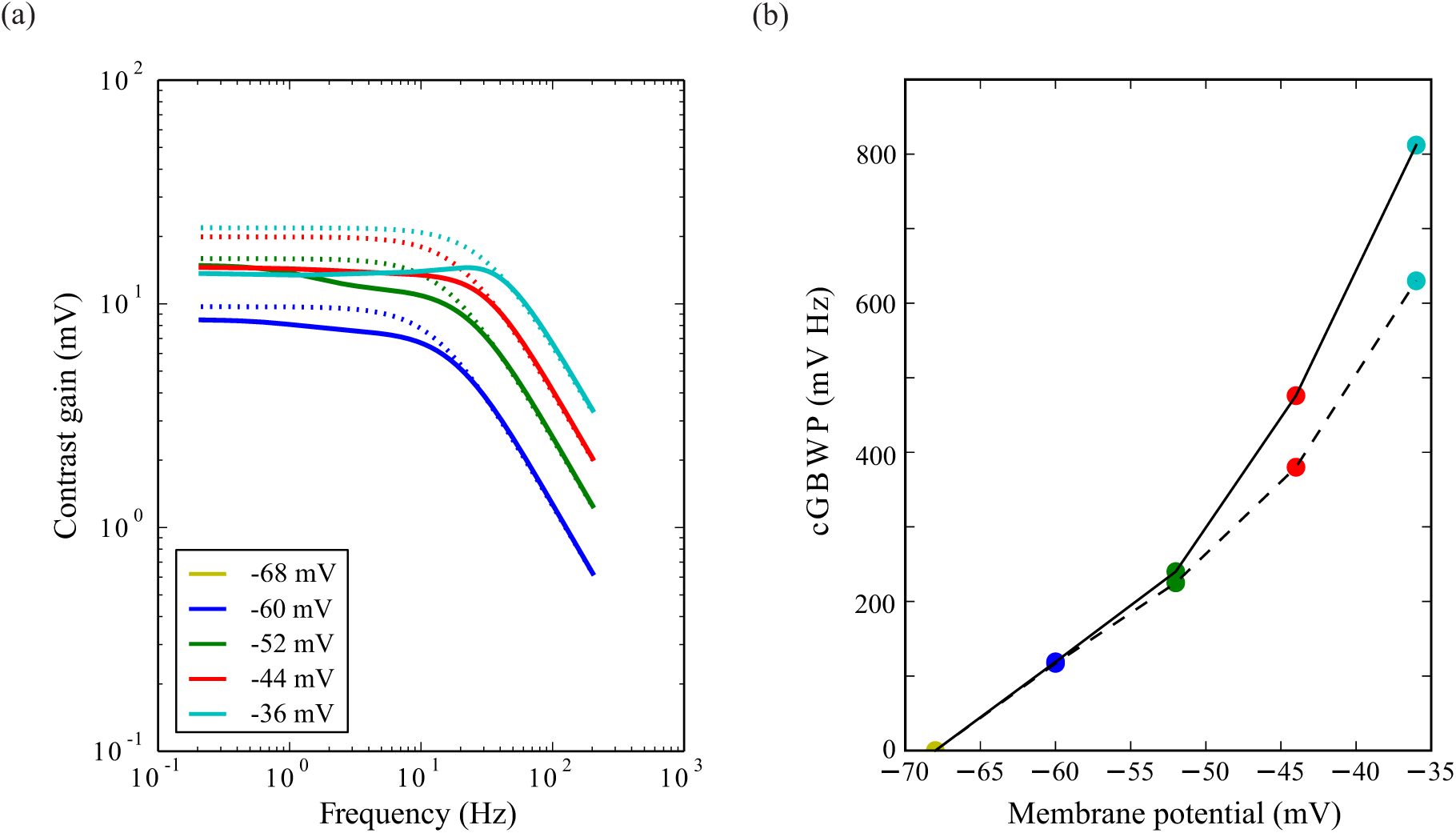
Voltage-dependent K^+^ conductances reduce contrast gain and increase contrast-gain-bandwidth product (cGBWP). (a) Contrast gain of the active photoreceptors (solid line) compared to the contrast gain of passive photoreceptors with the same resistance, depolarising leak conductance and capacitance (dotted line). Photoreceptors depolarised by light-induced current (LiC) to steady-state values of membrane potential indicated by coloured points in (b). Contrast gain for the completely dark adapted photoreceptor does not appear in the log-log plot as it is 0 mV. (b) cGBWP of the active photoreceptors (solid line) compared to the cGBWP of passive photoreceptors with the same resistance, depolarising leak conductance and capacitance (coloured points joined with dashed line).

The contrast gain-bandwidth product (cGBWP) is the product of the bandwidth and the peak contrast gain. In our simplified model, the contrast gain is the product between light induced current and membrane impedance (Eq 25), so the cGBWP is the product of GBWP and LiC. Consequently, the cGBWP of the active membrane exceeds that of the passive membrane by 10-15% at low light levels increasing to 25% at high light levels (Fig 8b).

### Light Dependent Modulation (LDM) of voltage-dependent K^+^ conductances change membrane impedance, gain and energy consumption

A light-dependent modulation (LDM) has been described in *Drosophila* R1-6 photoreceptors, arising from the action of PIP2 upon microvillar-bound Shab channels, which shifts their activation 10 mV towards negative potentials [26]. To determine the effect of the LDM at different depolarisations, we kept the leak and light conductances unchanged, allowing the membrane potential to change to a new steady-state value. The Na^+^/K^+^ pump and Na^+^/Ca^2+^ exchanger current change accordingly.

The shift in Shab voltage-activation range produced by PIP2 depletion hyperpolarises the photoreceptor and decreases the membrane impedance (Fig 9a). Consequently, the LDM speeds up responses by widening membrane bandwidth (Fig 9) [26], and causes the effects of Shaker become less conspicuous. The increase in bandwidth is small at low light levels becoming more pronounced at higher light intensities, when the membrane potential is above –55 mV and the Shab current is larger. The Shab mediated band-passing at high light intensities becomes more prominent, and bandwidth increases by up to 18 Hz (Fig 9b). As well as increasing bandwidth, LDM affects photoreceptor energy consumption. At low light levels, LDM has little effect upon photoreceptor energy consumption but at high light intensities energy consumption is increased by as much as ∼10% (Fig 9c).

**Figure 9:**
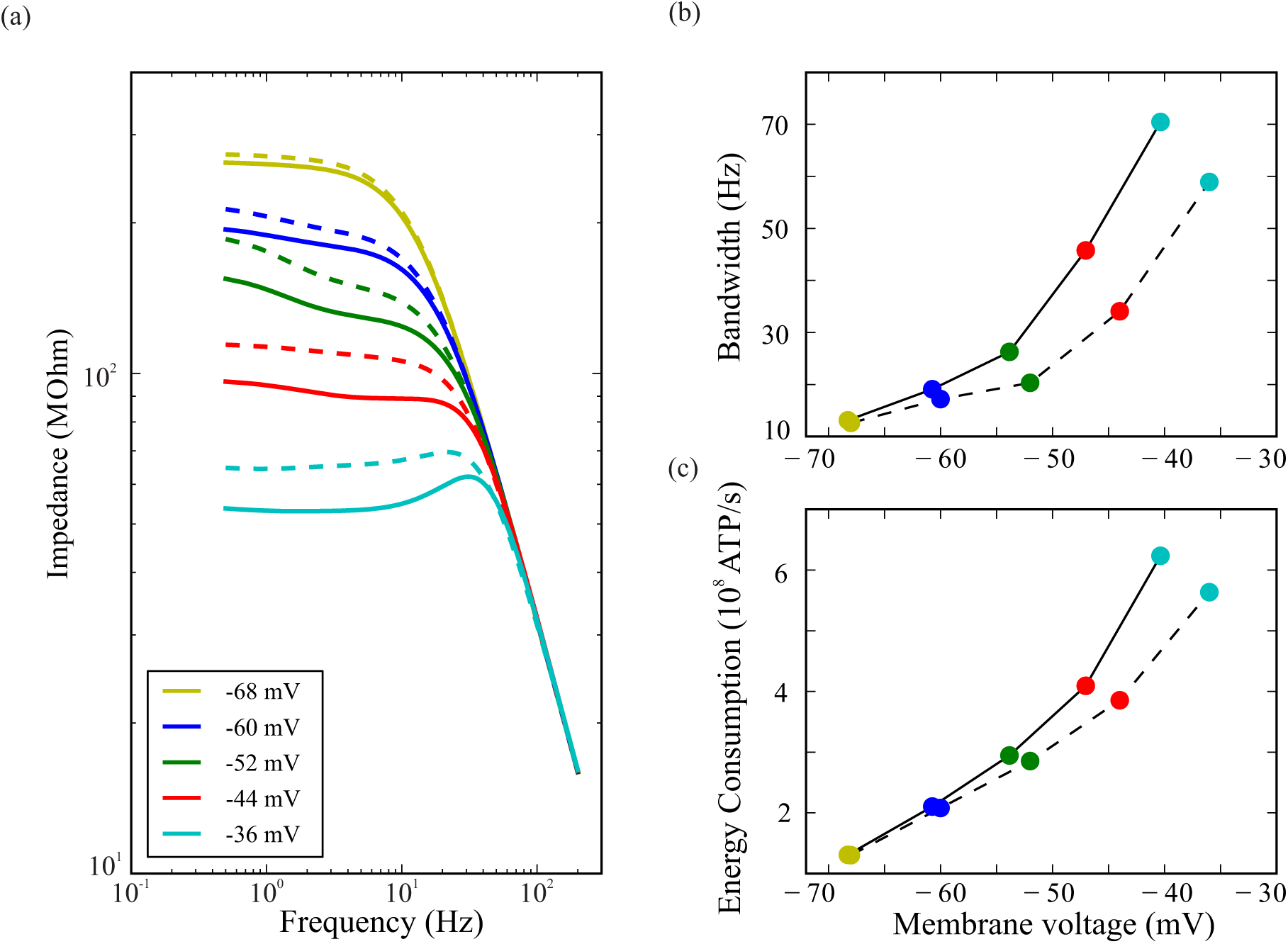
When PIP2 depletion shifts the voltage activation ranges of Shab, membrane impedance decreases, and bandwidth and energy costs increase. (a) Impedance with (solid lines) and without (dashed lines) the action of PIP2 depletion at four levels of depolarisation produced by changing the LiC. Dashed lines are the same as continuous lines in Fig 5a. (b) Bandwidth with (solid) and without (dashed) the action of PIP2 depletion. LiC fixed to values producing levels of depolarisation indicated by the key in (a). The leftward shift of coloured points indicates that PIP2 depletion hyperpolarises the membrane at rest, and at the 4 levels of LiC. (c) Energy cost, calculated as the rate of consumption of ATP, with (solid) and without (dashed) the action of PIP2 depletion on Shab.

Ca^2+^ activated calmodulin modulates the maximum conductance of Shab [52]. A decrease of 50% in the maximum Shab conductance hyperpolarises the photoreceptor and increases impedance, reducing the effect of Shab mediated band-passing at the highest light levels and reducing bandwidth by up to ∼20% (not shown). It also increases gain by ∼15% and decreases energy cost by up to ∼10%. Conversely, an increase of Shab maximum conductance, the expected effect of an increase in Ca^2^+/calmodulin upon light adaptation, has the opposite effect (not shown).

### Serotinergic modulation of voltage-dependent K^+^ conductances change membrane impedance, gain and energy consumption

Serotonin (5HT) reversibly shifts the activation and inactivation curves of the Shab conductance by ∼30 mV and the activation and inactivation curves of the Shaker conductance by 10 to 14 mV [24]. The higher activation voltages of the shifted K^+^ conductances caused by serotonin depolarises photoreceptors slightly and alters their filter properties (Fig 10). At each light-induced level of depolarisation photoreceptor impedance increases (Fig 10a) [25]. Serotonin also abolishes Shaker-mediated amplification by shifting inactivation [25]. Consistent with the reduction in the negative feedback contributed by Shab, serotonin decreases bandwidth at all depolarisations (Fig 10b), and by as much as ∼40% at high light levels. By modulating the properties of the Shab and Shaker conductances, serotonin also decreases energy consumption at all light levels by as much as ~40% (Fig 10c).

**Figure 10:**
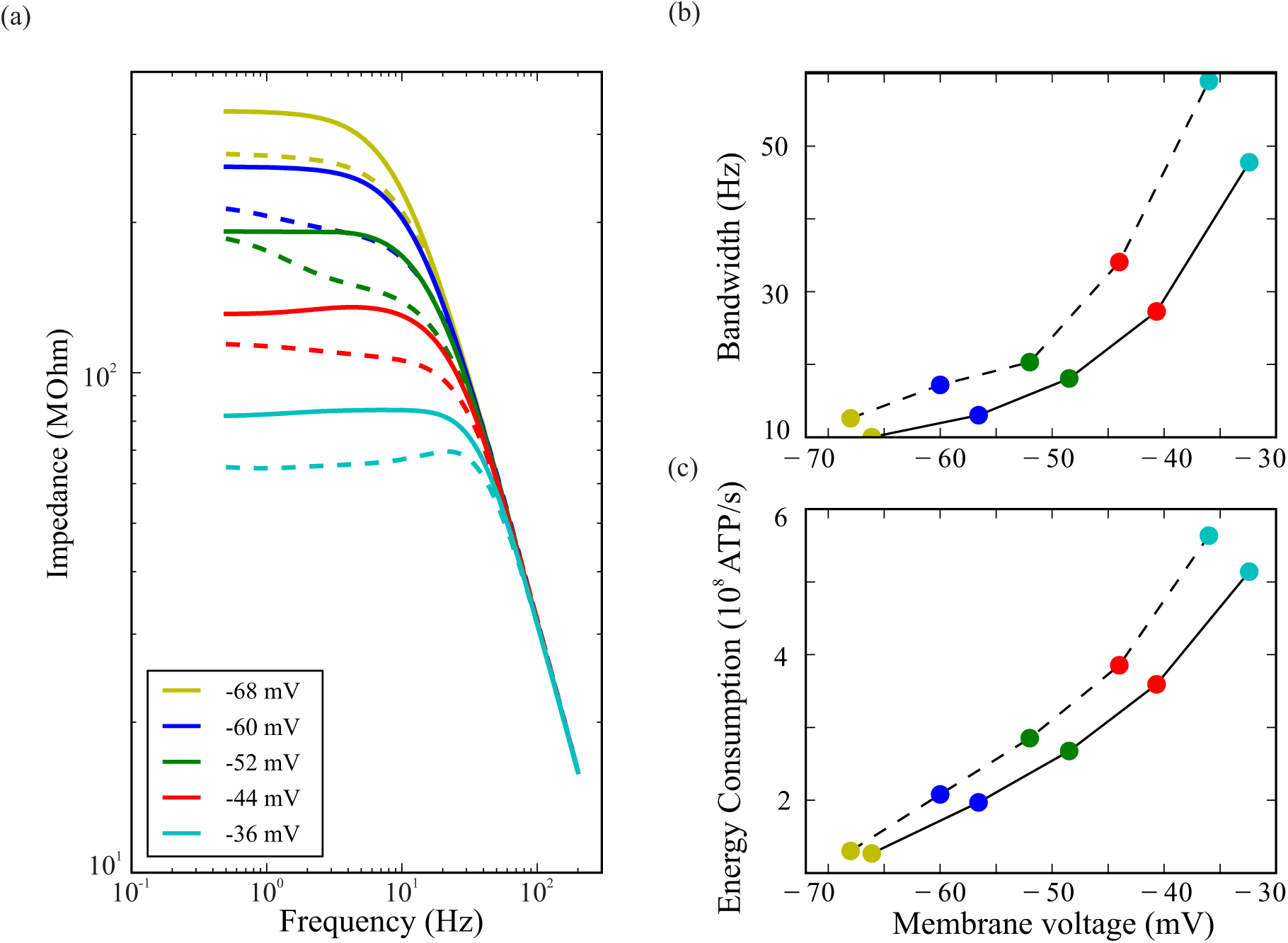
When serotonin shifts the voltage activation ranges of Shaker and Shab, membrane impedance increases, and bandwidth and energy costs decrease. (a) Impedance with (solid curves) and without (dashed curves) the action of serotonin at four levels of depolarisation produced by fixing the LiC. (b) Bandwidth with (solid) and without (dashed) the action of serotonin. LiC fixed to values producing levels of depolarisation indicated by the key in (a). The rightward shift of coloured points indicates that serotonin depolarises the membrane at rest, and at the 4 levels of LiC. (c) Energy cost, calculated as the rate of consumption of ATP, with (solid) and without (dashed) the action of serotonin on Shaker and Shab.

### Modulation of voltage-dependent K^+^ conductances adjusts the GBWP

By altering the voltage-dependent K^+^ conductances, modulation can adjust the gain-bandwidth product (GBWP) of the active photoreceptor membranes. Because the effects of modulation are specific, so too are the effects on shunt peaking and, consequently, the GBWP. Light-dependent modulation (LDM) has little effect at low and high light levels (−60 and −36 mV, respectively) but increases the GBWP at intermediate light levels (−52 to −44 mV) in comparison to the unmodulated active photoreceptor membrane (Fig 11a). The increase in GBWP is a consequence of LDM inducing a 10 mV negative shift in Shab activation so that it occurs at intermediate light levels producing shunt peaking, which extends bandwidth and increases the GBWP. Reduction of the Shab conductance by 50%, as produced by Ca^2+^ activated calmodulin modulation, reduces the photoreceptor GBWP at high light levels (−44 to −36 mV) but has no substantial effect at low and intermediate light levels (Fig 11b). The reduction in GBWP at high light levels is a consequence of the smaller Shab conductance, which reduces the shunt peaking produced by Shab activation and, consequently, the GBWP. Serotonin also reduces the GBWP at high light levels but additionally reduces the GBWP at low light levels and abolishes the ‘dip’ at intermediate voltages (−52 mV) producing a monotonic increase GBWP with depolarisation (Fig 11c). The effects on GBWP at low and intermediate light levels are due to the positive shift in the Shaker conductance caused by serotonin; Shaker activation at intermediate light levels causes shunt peaking that increases the GBWP, and simultaneously removes shunt peaking at low light levels. The positive shift in Shab conductance by serotonin reduces shunt peaking produced by Shab, thereby reducing the GBWP at high light levels.

**Figure 11:**
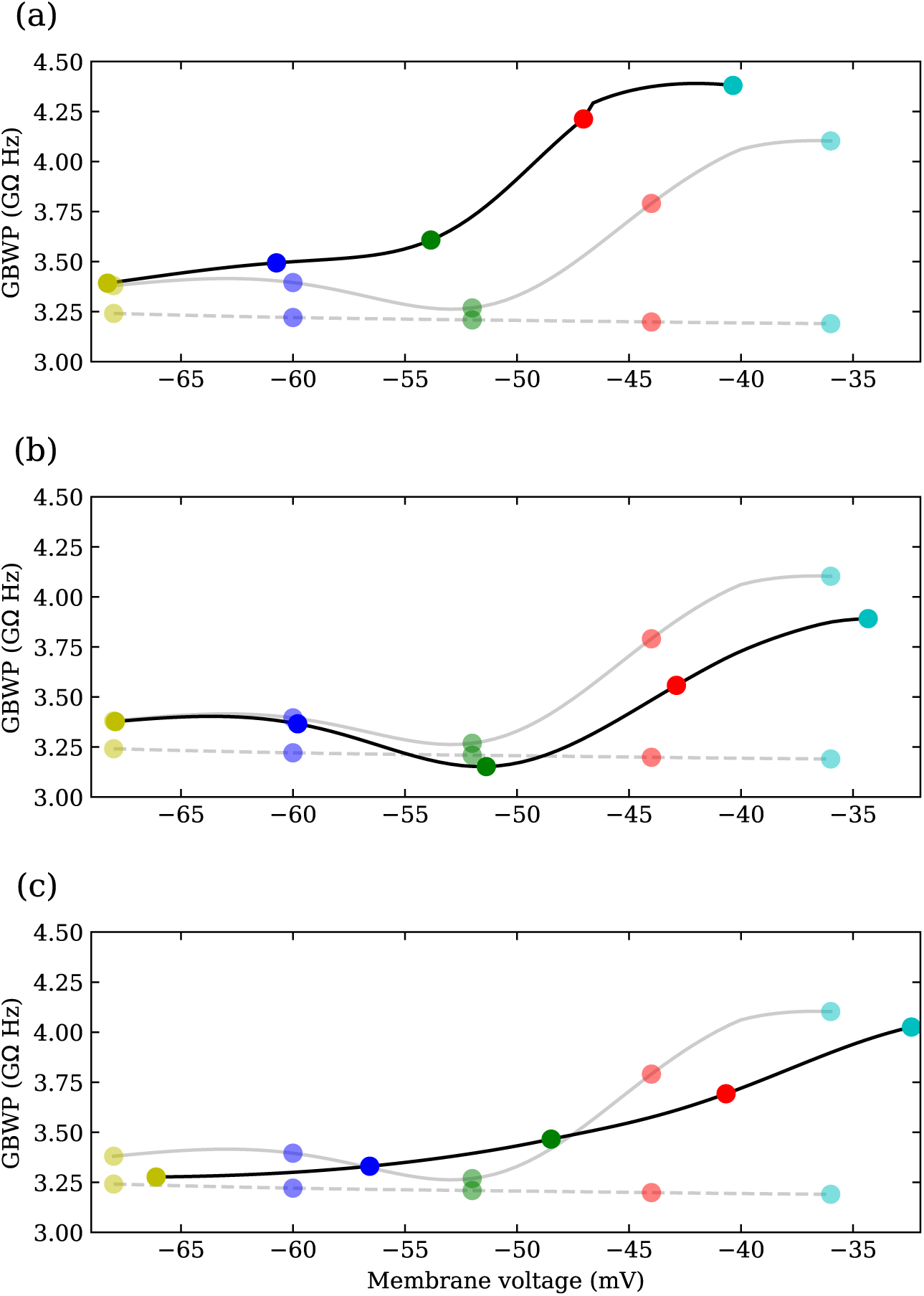
The contributions of different modulations to the gain-bandwidth product (GBWP) of the photoreceptor membrane, over the range of depolarisation produced by the light induced current. (a) Circles joined with black solid lines result from a model where Shab and Shaker have been modulated by PIP2 depletion. Circles joined with grey solid lines are the GBWP of the unmodulated photoreceptors, and circles joined with dashed lines are GBWP of the passive membrane, i.e. when all gating variables are frozen. (b) Same as (a) but reducing Shab conductance by 50%. (c) Same as (a) but with serotonin shifting Shaker and Shab activation and inactivation curves.

### Modulation of voltage-dependent K^+^ conductances trades-off gain for bandwidth but has little effect on the contrast gain bandwidth product (cGBWP) cost

The changes in photoreceptor membrane gain and bandwidth wrought by the modulation of voltage-dependent K^+^ conductances have implications for contrast coding. Light-dependent modulation (LDM) of the Shab conductance reduces photoreceptor contrast gain below the corner frequency at all light levels (Fig 12a), increasing photoreceptor bandwidth (Fig 13a). At the highest light levels, LDM enhances the bandpassing of the membrane. Conversely, serotonergic modulation of the Shab and Shaker conductances increases photoreceptor contrast gain below the corner frequency at all light levels (Fig 12b), decreasing photoreceptor bandwidth (Fig 13a). At the highest light levels, serotonin modulation removes the bandpassing of the membrane entirely.

**Figure 12:**
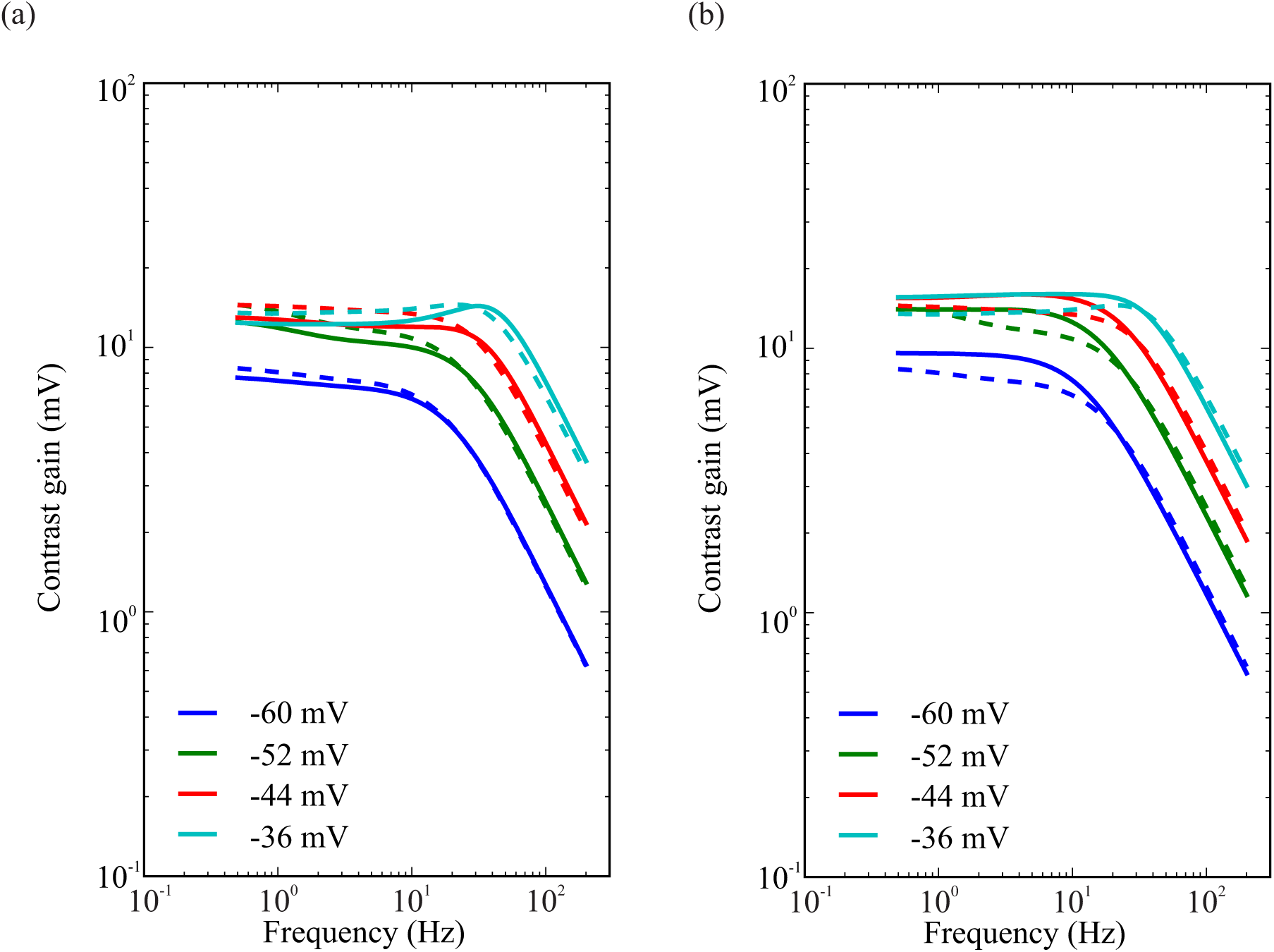
Contrast gain with (solid curves) and without (dashed curves) the modulator shift at four levels of depolarisation. (a) Light-dependent modulation (b) Serotonin.

**Figure 13:**
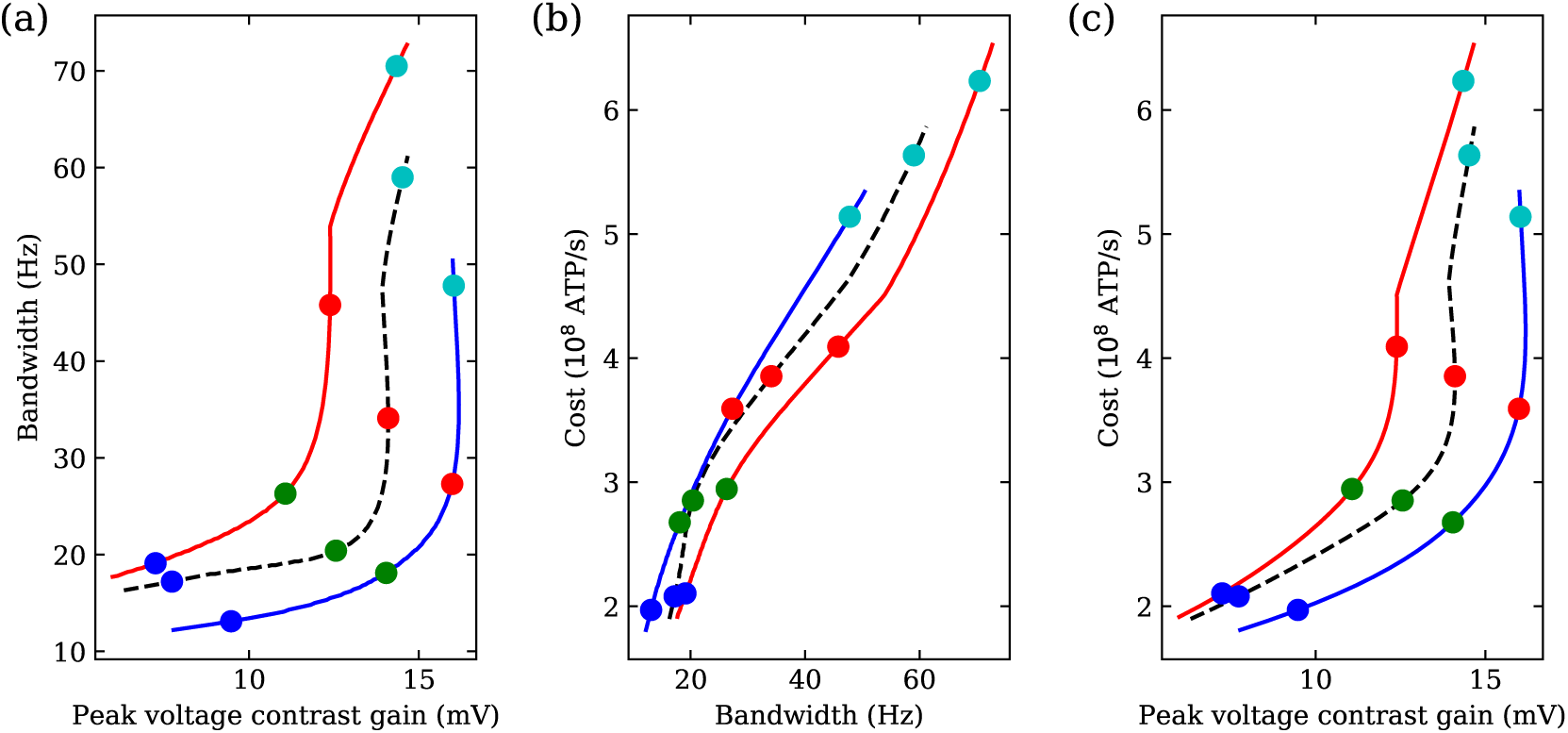
Modulators shift bandwidth, peak contrast gain and energy consumption of photoreceptors (a) Bandwidth and contrast gain plotted against each other with (solid curves) and without (dashed curves) modulation at four levels of depolarisation produced by fixing the LiC. Energy consumption and bandwidth plotted against each other with (solid curves) and without (dashed curves) modulation at four levels of depolarisation produced by fixing the LiC. (c) Energy consumption and peak contrast gain plotted against each other with (solid curves) and without (dashed curves) modulation at four levels of depolarisation produced by fixing the LiC.

To quantify the impact of modulation on the trade-off between contrast gain for bandwidth, we calculated the peak contrast gain and bandwidth at each light level (Fig 13). The dark adapted photoreceptor membrane has a low peak contrast gain and low bandwidth. At low light levels, peak contrast gain rises markedly though there is almost no change in bandwidth. At intermediate light levels, peak contrast gain rises further accompanied by a substantial increase in bandwidth, whilst at the highest light levels there is almost no change in peak contrast gain but bandwidth nearly doubles. Light-dependent modulation (LDM) shifts the relationship between peak contrast gain and bandwidth to higher bandwidths at an equivalent peak contrast gain. Conversely, serotonin modulation shifts the relationship between peak contrast gain and bandwidth to lower bandwidths at an equivalent peak contrast gain. Thus, modulation of the photoreceptor membrane is capable of trading-off contrast gain for bandwidth (Fig 14a).

**Figure 14:**
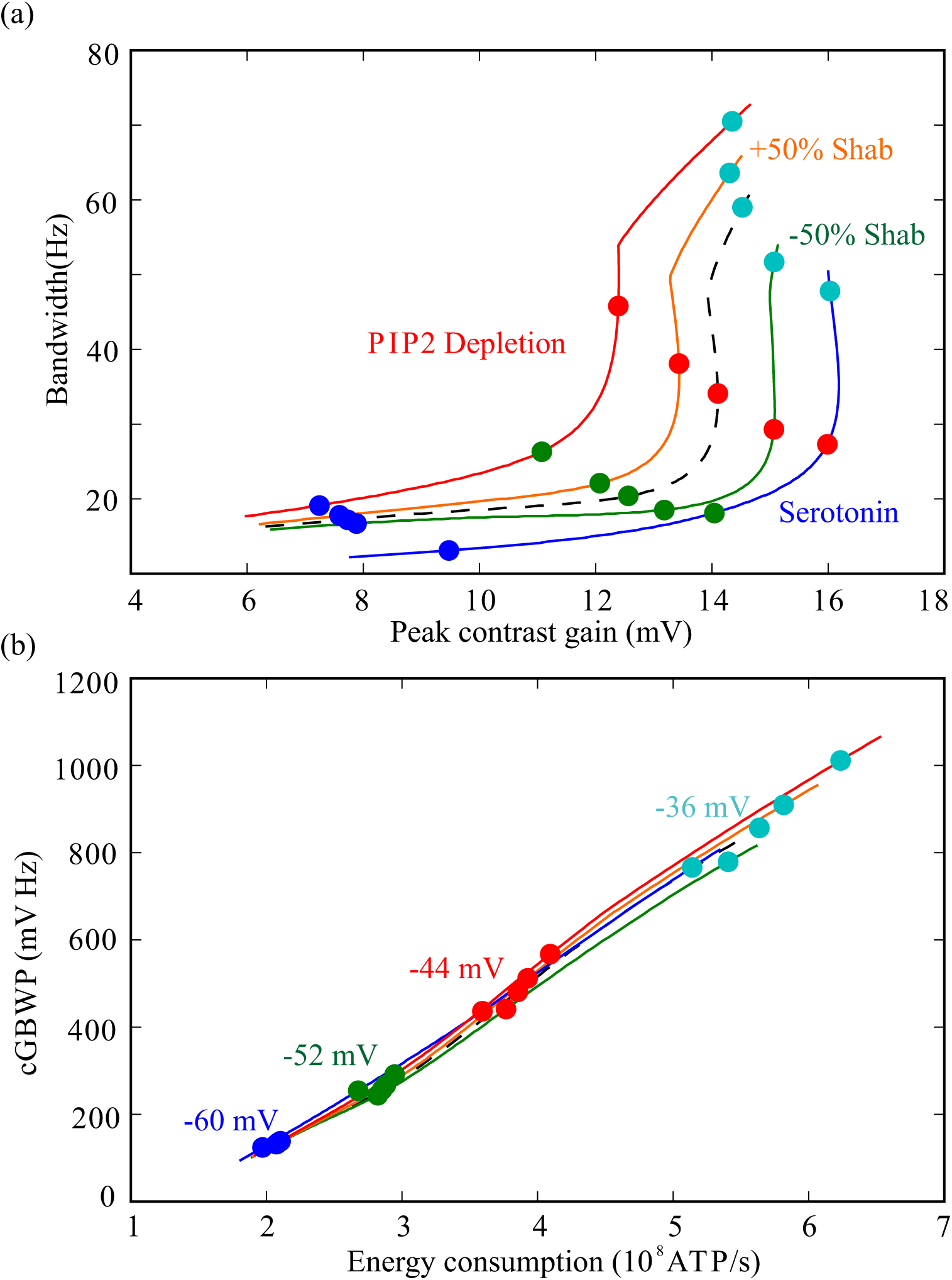
Shifts in voltage-dependent K^+^ conductances produced by modulators trade-off gain and bandwidth but have little influence on the cost of the contrast-gain-bandwidth product (cGBWP). (a) Bandwidth and peak contrast gain for the model photoreceptor with unshifted conductances (dashed black line) where light-induced conductance has been increased to depolarise to four membrane potentials (colour circles). At each of these membrane potentials the light-induced conductance remained constant, and the photoreceptor was allowed to reach a new depolarisation (coloured circles on coloured lines). (b) The original (black dashed line) and shifted (coloured lines, colour code as in (a)) photoreceptors occupy a similar space in a cGBWP vs energy cost graph, illustrating that modulators changing channel properties have little effect in the cost of cGBWP.

By altering photoreceptor energy consumption, modulation also alters the energy cost of bandwidth and of peak contrast gain. As light level increases, so too does bandwidth and energy cost. The energy cost of the peak contrast gain follows a similar relationship to that of bandwidth and peak contrast gain. Light-dependent modulation (LDM) reduces the energy cost of bandwidth but increases that of contrast gain (Figs 13b, 13c). Conversely, serotonin increases the energy cost of bandwidth and decreases that of contrast gain (Figs 13b, 13c). Thus, modulation adjusts the energy cost of peak contrast gain and bandwidth.

## Discussion

Our aim was to determine the impact of neuromodulators upon the energy efficiency of neural coding. To this end, we used a combination of computational and analytical models of the fruit fly R1-6 photoreceptor membrane that incorporate those voltage-dependent K^+^ conductances present in the soma. Our analysis demonstrated that the voltage-dependent K^+^ conductances fulfil previously unknown roles in photoreceptor coding, extending bandwidth, increasing the GBWP and improving energy efficiency. Using these models, we assessed the effect that changes in the voltage-dependent K^+^ conductances produced by the action of neuromodulators have upon membrane filtering, signalling and energy efficiency. We show that neuromodulators trade-off contrast gain for bandwidth but do so without producing a substantial change in energy consumption, so that the energy cost of cGBWP remains unchanged (Fig 14b). Instead, modulation acts to adjust the energy investment in contrast gain versus bandwidth whilst maintaining the relationship between cost and cGBWP.

The voltage-dependent K^+^ channels present in fly photoreceptors reduce gain and increase bandwidth by reducing membrane resistance upon depolarisation [53, 18] and by reducing and shaping the impedance [23, 14]. By altering the properties of the voltage-dependent K^+^ channels, neuromodulators directly affect photoreceptor performance. Serotonin increases *D. melanogaster* photoreceptor gain increasing the amplitude of signals at low light intensities, potentially enhancing the ability of the visual system to detect events. Simultaneously, serotonin reduces investment in high bandwidths, which are absent at low light intensities thereby saving energy. Conversely, PIP2 reduces gain at high light intensities when signals are large and easily detected, saving energy. Instead, PIP2 increases bandwidth at high light levels enabling the visual system to detect faster signals. By acting on the first layer of the visual system, and through their potential concerted effects upon other cell types, these modulators have the potential to affect processing throughout the visual system.

### Assessing performance using the gain-bandwidth product (GBWP)

We assessed photoreceptor membrane performance by calculating the product of bandwidth and the peak of the impedance function, the gain-bandwidth product (GBWP). This measure combines two biologically germane features of the membrane, gain and bandwidth; a sufficiently high gain can prevent signals from being corrupted by noise, whilst a sufficiently high bandwidth permits fast signals to be transmitted. In both cell membranes and operational amplifiers, the GBWP determines the maximum gain for a given bandwidth, and vice versa. In cell membranes, the GBWP is determined by the membrane capacitance, and specific membrane capacitance is relatively invariant. When ion channels reduce the membrane resistance or produce negative feedback, gain is traded for bandwidth. Only by modifying the one-pole filtering of the membrane (e.g. by using shunt-peaking) can the membrane increase the GBWP and thus increase gain for a given bandwidth or bandwidth for a given gain. Thus, the GBWP provides a single variable allowing quantification of the overall photoreceptor membrane performance. We extended this idea to include the light-induced current (LiC). A useful concept when describing the photoreceptor response to light is the contrast gain, the frequency-dependent gain between light contrast and voltage in the photoreceptor soma. As the product of the LiC and the membrane impedance, the contrast gain combines the outputs of the phototransduction cascade and the photoinsensitive membrane. The contrast gain-bandwidth product (cGBWP), is the product of the bandwidth and the peak contrast gain, quantifying the impact of the membrane upon contrast gain and bandwidth. As the cGBWP is also the product between GBWP and LiC, it can be increased partially through shunt-peaking, but also through an increase in LiC at more depolarised membrane potentials.

### Voltage-dependent K^+^ conductances generate negative feedback and shunt peaking that extend bandwidth and reduce cost

We began by assessing the impact of voltage-dependent K^+^ conductances themselves upon the graded electrical signalling and energy consumption of fruit fly photoreceptor membranes. By activating upon depolarisation, the Shaker, Shab and Novel conductances produce negative feedback. The main role of this feedback is to reduce the impedance to below that of a passive photoreceptor with identical membrane resistance and capacitance, as is the case for delayed rectifier conductances in blowflies [14]. Activation of the Shaker conductance at low depolarizations and the Shab conductance at high depolarizations produces shunt peaking, which is again produced by delayed rectifier conductances in blowflies [23]. Shunt peaking is produced by voltage-dependent ion channels changing their conductance with membrane potential after a delay, causing them to act like an inductance. This increases bandwidth without decreasing membrane gain, thereby increasing the photoreceptor membrane gain-bandwidth product (GBWP). The combination of the decrease in gain by negative feedback and the increase in GBWP by shunt-peaking caused by voltage-dependent K^+^ conductances produces an increase in bandwidth without incurring the full cost that reducing the membrane resistance of a passive membrane would produce. Consequently, voltage-dependent conductances modify the trade-offs between membrane bandwidth, gain and energy cost by making it cheaper to operate a membrane with lower gain and higher bandwidth.

### Shaker and Shab inactivation generates positive feedback amplifying low frequencies and saving energy

At the resting potential and at low light levels a window current [40] is present in *D. melanogaster* photoreceptors produced by the overlap between the activation and inactivation of the Shaker K^+^ conductances [22, 49, 25]. The reduction in the window current upon depolarisation has been assumed by previous studies to produce amplification at low frequencies in *D. melanogaster* photoreceptors [22, 49, 25]. Our biophysical models show that this increase in the gain of the membrane at low frequencies occurs against a background of negative feedback produced by the activation of other conductances as well as that of Shaker itself. In some *D. melanogaster* photoreceptors, amplification may be sufficiently prominent so as to produce an impedance that at low frequencies is above the membrane resistance (Frolov, pers. comm.). However, similarly to our model, in most photoreceptors the impedance is unlikely to exceed the membrane resistance, and so it is rather a reduction in attenuation.

Amplification by the Shab conductance at high light levels has not, to our knowledge, been described before. The amplification is restricted to lower frequencies than that of Shaker, because of the slower inactivation dynamics of Shab. It is smaller because Shab inactivation occurs at high light levels when input resistance is low, reducing the relative effect of feedback. Moreover, the Shab steady-state inactivation curve has a relatively shallow slope and is centred at –25.7 mV, outside the range of typical steady-state depolarisations.

The functional significance of low frequency amplification remains unclear. Indeed, low frequency attenuation by the light current [37] makes it likely that the amplification produced by the Shaker and Shab conductances has little or no impact upon signalling in fruit fly photoreceptors, and is merely a side effect of inactivation. Thus, the role of Shaker in fruit fly photoreceptors is similar to that of A-type conductances in the dendrites of vertebrate interneurons: decreasing the amplitude and time-to-peak of depolarising postsynaptic potentials rather than amplifying them [54]. Our modelling suggests that by increasing bandwidth at low light levels whilst ensuring that high bandwidths do not occur at high light levels, Shaker conductance inactivation reduces energy consumption in comparison to an equivalent non-inactivating conductance (Fig 7), as hypothesised by Niven *et al.* [21]. This raises the possibility that the inactivation of A-type conductances in dendrites [54] may also be linked to reducing energy consumption.

### Feedback from molecular exchangers and pumps

Other molecular components of the photoreceptors can also produce low frequency amplification. For example, the Na^+^/Ca^2^+, which has a time constant of 350 ms [34], can produce positive feedback amplifying frequencies below 1 Hz. However, the fraction of Ca^2^+ in the LiC is only about 26% [34], and thus the exchanger net current is only 13% of the LiC, making the contribution of this positive feedback small. Moreover, this positive feedback is unlikely to affect photoreceptor bandwidth and gain because the light response filters low frequencies [37]. To maintain consistency with other choices of our model, such as only considering frequencies above 2 Hz in bandwidth and gain calculations, we considered the Na^+^/Ca^2^+ exchanger as a constant current at each steady-state depolarisation, while studying the response to small signals.

Our models also incorporated the Na^+^/K^+^ pump as a constant current at each steady-state depolarisation, rather than incorporating the actual dynamics. This approximation is accurate for the Na^+^/K^+^ pump current because its slow dynamics (seconds, e.g. [29]) restricts its negative feedback to extremely low frequencies.

### Modulation of voltage-dependent K^+^ conductances adjusts investment in photoreceptor gain and bandwidth

Neuromodulators alter specific properties of the voltage-dependent K^+^ conductances, shifting the trade-off between contrast gain and bandwidth in fruit fly photoreceptors. Light-dependent modulation (LDM) of the Shab conductance by PIP2 depletion, which shifts Shab activation to lower potentials [26], moves the trade-off to favour higher bandwidths and lower peak contrast gains. Conversely, the positive shift in the voltage operating range of both the Shaker and Shab conductances caused by serotonin [24], moves the trade-off to favour lower bandwidths and higher peak contrast gains. Likewise, Ca^2^+/Calmodulin antagonists that decrease the Shab conductance [52] also favour lower bandwidths and higher peak contrast gains. Conversely, an increase in the Shab conductance would favour higher bandwidths and lower peak contrast gains. So, modulators enable fruit fly photoreceptors to explore a greater region of the trade-off between peak contrast gain and bandwidth. This enables photoreceptors to adjust gain or bandwidth depending on the prevailing conditions. Both shifts produced by products of the photocascade, PIP2 depletion and an increase in Shab conductance produced by an increase in Ca^2^+ [26, 52], sacrifice gain for bandwidth thereby contributing to light adaptation (Fig 14a, 15). Serotonin [24] has the opposite effect, increasing gain at the cost of bandwidth potentially adjusting photoreceptors to low light levels (Fig 14a, 15).

**Figure 15:**
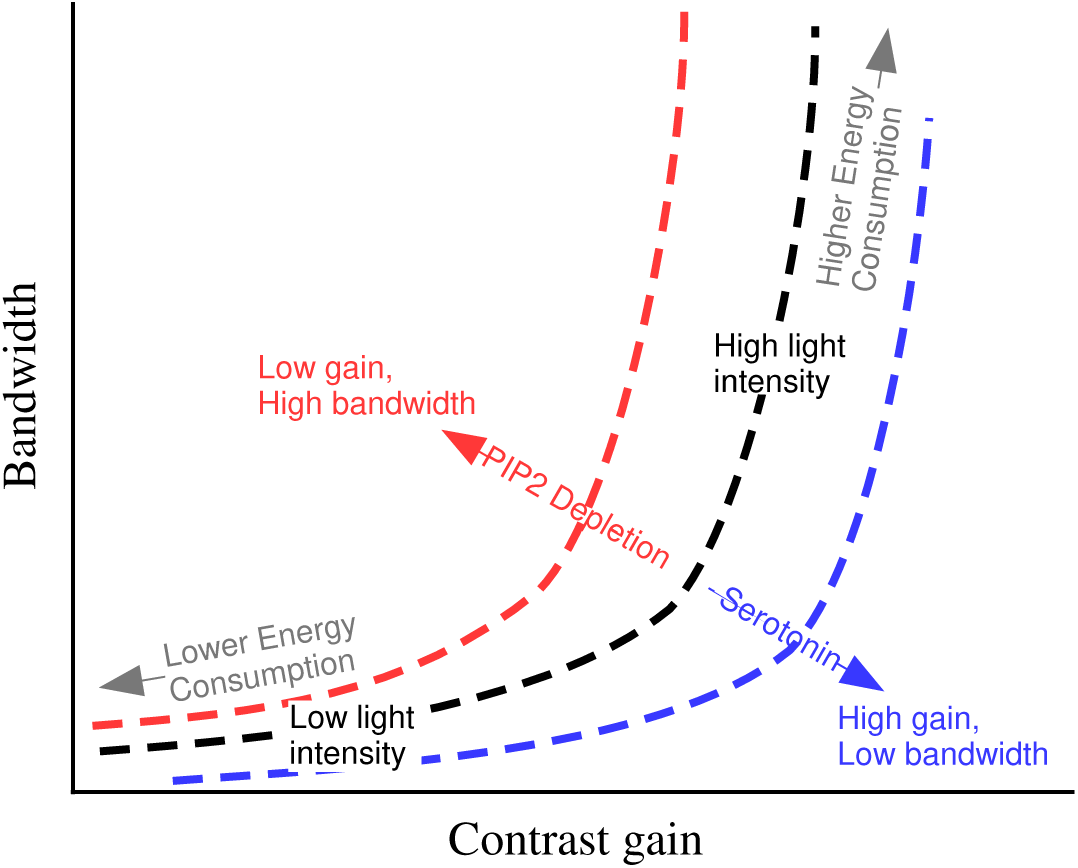
Neuromodulators trade-off photoreceptor gain and bandwidth. The model photoreceptor is shown with unshifted conductances (dashed black line) or with conductances shifted by neuromodulators (coloured lines).

By adjusting voltage-dependent conductances, these neuromodulatory changes also have implications for photoreceptor energy consumption. If the light-induced conductance is kept constant, the neuromodulatory effects of phototransduction cascade products [26, 52] that increase bandwidth also increase in energy consumption. Thus, they reinforce the ability of the membrane to produce high bandwidth only when it is needed. This allows photoreceptors to save energy by avoiding investing in high bandwidth when it is not needed. Conversely, serotonin [24] reduces bandwidth and decreases the energy consumption if the light-induced conductance is kept constant. For a given energy consumption, light-dependent modulation (LDM) by PIP2 depletion decreases the cost of bandwidth but increases the cost of gain (Fig 13b,c). Conversely, serotonin decreases photoreceptor bandwidth but increases gain (Fig 13b,c). Likewise, Ca^2^+/Calmodulin antagonists also increase the energy cost of bandwidth and decreases the energy cost of gain. An increase in Ca^2^+/Calmodulin also decreases the cost of bandwidth but increases the cost of gain. The presence of multiple voltage-dependent conductances combined with multiple neuromodulators allows the costs of gain and bandwidth to be adjusted in independent ways.

By shifting the balance between gain and bandwidth (Fig 14a) neuromodulators can change the contrast-gain-bandwidth product (cGBWP). However, they have little effect on the cost of the cGBWP (Fig 14b). This can be understood by considering that the cGBWP is the product of the membrane GBWP times the light current, while the energy cost is proportional to the depolarising current —which is dominated by the light current at all but the lowest light levels. Thus, the cost of cGBWP is proportional to the membrane GBWP. As GBWP tends to be restricted between 1.2 and 1.5 for most choices of conductances, the cost of cGBWP varies little upon depolarisation or conductance shifts.

### Implications of the simplified model of the light-induced current

We used a simplified model of phototransduction in which the photocurrent follows photon absorption. This model fails to capture low frequency filtering by light adaptation and high frequency filtering by the finite width and latency dispersion of the quantum bump [44, 45]. Indeed, in *Drosophila* photoreceptors the phototransduction cascade current has a bandwidth between two and five times lower than the membrane bandwidth [37].

Incorporating the light-induced current (LiC) into the model allowed us to study photoreceptor gain in a more relevant way than with membrane impedance alone. If membrane impedance is decreased using negative feedback, no increase the LiC is needed to keep the membrane voltage steady. The decreased impedance reduces the membrane gain but, because the LiC is unchanged, the net result is a decrease in contrast gain. Conversely, if the membrane impedance were decreased by reducing the membrane resistance, an increase in the LiC would be necessary to keep the membrane voltage steady. The net result would be that contrast gain is unchanged. This illustrates the importance of the interplay between the LiC and the membrane impedance.

### Implications for arthropod photoreceptors

Within the arthropods, neuromodulation of photoreceptors is widespread. Some of these neuromodulators target the voltage-dependent conductances, whereas others act on molecular components of photoreceptor phototransduction cascade. In the fruit fly (*Drosophila melanogaster*), both octopamine and dopamine can increase the latency of quantum bumps without changing the membrane properties [55].

Likewise, in the desert locust (*Schistocerca gregaria*) serotonin shifts photoreceptors from the day state in which they express a voltage-dependent K^+^ conductance with delayed rectifier-like properties to the night state in which they express a rapidly inactivating voltage-dependent K^+^ conductance [56]. Just as in *D. melanogaster* photoreceptors, the modulation of voltage-dependent K^+^ conductances by serotonin in desert locusts is likely to boost gain at low light intensities, increasing quantum bump amplitude thereby making them easier to detect. Increasing the expression of a rapidly inactivating voltage-dependent K^+^ conductance will also likely reduce locust photoreceptor energy investment in bandwidth, producing a similar redistribution of energy investment in gain and bandwidth that our modelling shows is produced by serotonergic modulation in *D. melanogaster* photoreceptors. The light response of desert locust photoreceptors is also increased by nitric oxide and cGMP [57], again improving detectability.

The combination of neuromodulators that adjust elements of the phototransduction cascade and the photoinsensitive membrane separately provides considerable flexibily within photoreceptors to match prevailing environmental conditions, thereby ensuring that energy is invested in high gains or high bandwidths only when it is needed. Avoiding expenditure on high bandwidth when it is unnecessary is an important strategy for reducing the cost of vision. Such changes may be predicted rather than being generated by prevailing environmental conditions, such as in the horseshoe crab (*Limulus*) in which photoreceptors undergo 24-hour cyclic changes in sensitivity caused by octopamine (for a review see [58]).

### Implications for neuromodulation

The effects of neuromodulators upon the dynamics of single neurons and neural circuits are well documented in both invertebrates and vertebrates [1, 2, 59]. Whilst neuromodulatory effects on the performance of single neurons and neural circuits have been described extensively, their impact upon the energy consumption has been largely ignored (but see [60, 61]). Our results show that the changes wrought by neuromodulators to voltage-dependent conductances are likely to affect the energy consumption of neurons and neural circuits. Indeed, many changes caused by neuromodulators, such as to sensory or synaptic inputs/outputs, or the reconfiguration of neural circuits, have implications for neuronal energy consumption. Although some changes may increase or decrease overall energy consumption [60, 61], our findings suggest that the impact may be more subtle; neuromodulators may redistribute energy investment in different aspects of neuronal performance without substantially altering overall energy consumption.

Mechanoreceptors demonstrate the potential for neuromodulators to adjust or redistribute neuronal energy consumption. Neuromodulators can adjust numerous properties of invertebrate mechanoreceptors including their firing rate, dynamics, sensitivity and spike timing precision [62, 63, 64, 65, 66, 67, 68, 69, 7]. Although in many cases the underlying causes of these changes are unknown, in some they are linked to changes in conductance. For example, in the crab (*Carcinus maenas*) neuromodulation by allatostatin or serotonin of mechanoreceptors can adjust their spike rates and spike timing precision through changes in conductance; high conductance states are associated with lower spike rates and higher spike timing precision [7]. In such cases, there will be implications both for the energy consumption of the neuron and for the cost of information coding. High conductance states to increase spike timing precision caused by allatostatin will likely increase the energy consumption of spikes and of maintaining the resting potential but the accompanying reduction in spike rate may reduce energy consumption [7]. In these mechanoreceptors, serotonin has the opposite effect producing a low conductance, high spike rate state. Thus, as in fruit fly photoreceptors, neuromodulators in crab mechanoreceptors may redistribute the energy invested in different aspects of neuronal coding.

The ability of neuromodulators to redistribute the energy invested in different aspects of neuronal performance, and potentially to adjust the overall energy expenditure, has implications not only for single neurons but also for neural circuits. There are potential energy savings to the reconfiguration of neural circuits by neuromodulators to generate different behavioural outputs. The reuse of neurons to generate different behaviours reduces the need to build and maintain additional neurons, which would reduce energy consumption [70]. Neuromodulators may also redistribute energy investment in different parts of neural circuits. Indeed, neuromodulation may allow concerted changes in neurons and across neural circuits to co-ordinate investment in different aspects of neural processing to match prevailing behavioural and environmental circumstances.

### Supporting information

## Acknowledgments

The authors thank SB Laughlin, M Lengyel and DG Stavenga for discussion and feedback on previous versions of the manuscript.

FJHH was supported by Fundación Caja Madrid, The Department of Zoology of the University of Cambridge and Trinity College. JEN was supported by the BBSRC (grant number BB/R005036/1).

